# Basigin Links Altered Skeletal Stem Cell Lineage Dynamics with Glucocorticoid-induced Bone Loss and Impaired Angiogenesis

**DOI:** 10.1101/2025.01.25.634887

**Authors:** Thomas H. Ambrosi, David Morales, Kun Chen, Ethan Hunt, Kelly Weldon, Amber N. Maifeld, Fatima I.M. Chavez, Yuting Wang, Matthew P. Murphy, Amin Cressman, Erika E. Wheeler, Augustine Saiz, J. Kent Leach, Fernando Fierro, Charles K.F. Chan, Nancy E. Lane

**Affiliations:** Department of Orthopaedic Surgery, University of California at Davis Medical School, Sacramento, CA, USA; Stem Cell Institute, Stanford University Medical School, Stanford, CA, USA; Institute for Regenerative Cures, University of California Davis, Sacramento, CA, USA; Department of Biomedical Engineering, University of California at Davis, Davis, CA, USA; Department of Medicine, Division of Rheumatology, University of California, Davis, Sacramento, CA, USA

## Abstract

Glucocorticoid (GC) induced osteoporosis (GIOP) and osteonecrosis remain a significant health issue with few approved therapies that can treat the bone loss and dysfunction of skeletal vasculature. Therefore, we aimed to investigate the cellular and molecular processes by which GCs affect osteogenesis and angiogenesis, as well as how treatment with parathyroid hormone (hPTH 1-34) modifies these effects in a mouse model of GIOP. GC treatment reduced bone mass through decreased bone formation by skeletal stem cells (SSCs) while also increasing osteoclast mediated resorption. Concomitantly, endothelial cells were increased in numbers but displayed distorted phenotypical features. However, hPTH treatment reversed GC induced changes in osteogenesis and angiogenesis to control levels. Transplantation studies of SSCs combined with molecular analysis by single cell RNA-sequencing and functional testing of primary human cells tied GC-induced skeletal changes to altered stem and progenitor cell differentiation dynamics. This in turn perpetuated reduced osteogenesis and vascular malformation through direct SSC-endothelial crosstalk mediated at least in part by Basigin. Intriguingly, antibody-mediated blockade of Basigin during GC treatment prevented detrimental bone loss. In addition, when administered to aged mice, anti-Basigin therapy reinstated bone remodeling to significantly improve bone mass independent of sex. These findings, while helping to explain the cellular and molecular basis of how hPTH treatment can mitigate GC induced bone loss, provide new therapeutic vantage points for GIOP and other conditions associated with bone loss.

## Introduction

Glucocorticoids (GCs) are potent anti-inflammatory compounds, however, continued exposure results in bone loss and osteonecrosis^1^. Since SSCs are crucial for maintaining skeletal homeostasis, it is tempting to speculate that GCs might affect their function. Recently, detrimental changes to angiogenesis have also been implicated in GC-induced bone loss^2,3^. Nevertheless, the mechanism by which GCs alter vascularity remains unknown. Our study aimed to evaluate GC-effects on both osteogenesis and angiogenesis, as well as if treatment with parathyroid hormone (hPTH 1-34) modifies the effects in a mouse model of GC induced bone loss.

GCs are frequently prescribed for both acute and chronic medical conditions to reduce inflammation and immune system activity. GCs are known to alter bone remodeling by reducing osteoblast activity, stimulating osteoclast maturation and activity and reducing gonadal hormone production which indirectly activates osteoclastogenesis^1^. Bone loss induced by GCs tends to have two phases, with an initial rapid loss of bone mass, particularly trabecular, followed by a sustained gradual reduction that results in the loss of cortical bone. Independent of the GC-induced loss of bone mass, subjects treated with GCs often rapidly reduce mechanical strength of bone such that many subjects experience fractures while treated^4^.

After the initial stages of GC-induced bone loss, a continued suppression of bone formation has been reported. This is mainly due to reduced activity of osteoblasts, the differentiated cells that are responsible for bone formation derived from a pool of SSCs. hPTH 1-34, also known as teriparatide, is FDA-approved to treat GC-induced osteoporosis as it stimulates osteoblasts to generate bone in the presence of GCs^5,6^, decreases sclerostin production by osteocytes and prevents osteoclast maturation in the presence of GCs^7^.

GC exposure also can induce osteonecrosis, the result of damaged skeletal vasculature that provides essential nutrients for normal cell function and tissue homeostasis. This adverse event is commonly associated with higher cumulative doses and longer treatment courses of systemic GCs, however. it has also been described after intra-articular injections, topical administration or low-dose, short-term oral steroids^3^. No major gender differences have been identified. While the exact etiology of osteonecrosis is not clear, reduction in the vascular supply to the proximal femur is frequently observed. Angiogenesis is required for the osteogenic process, as endothelial cell invasion is required to transition hypertrophic chondrocytes to osteoblasts at the growth plate, and angiogenesis through macrophage initiated signaling is needed for the formation of a bone remodeling unit. However, the influence of GCs on angiogenesis is unclear^8^.

Therefore, given the observation that GCs reduce bone formation and may alter vascular supply directly or indirectly to bone, the purpose of this study was to increase our understanding of how GCs affect the cellular and molecular components of the bone marrow environment. We performed a detailed analysis of the cellular composition, SSC activity and single cell gene expression in mice treated with GCs, after recovery from GCs and GC with hPTH 1-34 treatment. We determined that bone loss through GC exposure is initially associated with a reduced number of SSCs and vascular progenitor cells, while with continued GC exposure skeletal precursor cells accumulated but lost their ability to differentiate into osteogenic and chondrogenic cell types. Interestingly, these SSCs secrete specific factors, including Basigin, that alter endothelial morphology and function through direct crosstalk. Concomitant hPTH 1-34 during GC exposure or antibody-mediated Basigin blockade abrogated that detrimental signaling axis and reversed the GC-mediated phenotype. Excitingly, anti-Basigin treatment also improved bone parameters in aged mice suggesting a potential broader clinical utility. Altogether, these findings reveal a previously unappreciated connection between stem and endothelial cell interaction that can be targeted to prevent GC-induced bone loss.

## Results

### GC-induced bone loss is reversed by concurrent hPTH treatment

To assess the cellular and molecular changes of continuous GC exposure on bone tissue, we subcutaneously transplanted 4-month-old, male Balb/cJ mice with 5 mg Methylprednisolone (GC) or placebo release pellets. Randomly grouped mice were housed for 28 days before analysis. To assess recovery from GC treatment, additional animals were kept for another 28 days with separate groups having GC pellets removed and animals receiving hPTH 1-34 for the second half of the experiment at 40 μg/kg, 5 days/week with GC exposure (Fig.1a). As expected from our previous study using a similar model^9^, micro-CT analyses and mechanical testing showed that 28- and 56-day GC exposure significantly reduced femoral trabecular bone parameters and bone strength, respectively, compared to placebo control (Fig.1b-d & Extended Data Figure 1a-b). Interestingly, removing GC pellets after 28 days did not reverse trabecular bone loss after an additional 28 days, while hPTH 1-34 in the presence of GC exposure normalized trabecular parameters to placebo levels. Cortical thickness and area were significantly reduced upon acute GC exposure but were unaltered in all groups monitored over a 56-day period (Extended Data Figure 1c-d). To test how these observations were tied to changes in bone remodeling, we conducted dynamic histomorphometry and found that bone formation and osteoclast activity were inversely controlled by GCs. Mineral apposition and bone formation rate were significantly reduced by GC treatment at both investigated timepoints and only hPTH 1-34 treatment improved those parameters to placebo control levels (Extended Data Figure 2a-c). In contrast, osteoclast activity was strongly increased upon GC exposure as measured by TRAP staining and bone marrow-derived in vitro osteoclastogenesis; however GC removal alone was sufficient to reduce bone resorption activity to control levels (Extended Data Figure 2c-d). Osteoclast surface per bone surface was elevated in GC mice that received hPTH 1-34 suggesting stimulation of increased bone remodeling with high bone formation rates outweighing increased bone resorption. Of note, the growth plate height was initially reduced upon GC stimulation but normalized to placebo levels at day 56 (Extended Data Figure 1e). Altogether, these data provide insight into the complex effects of GC and hPTH 1-34 actions on bone parameters of our mouse model.

**Figure 1.**
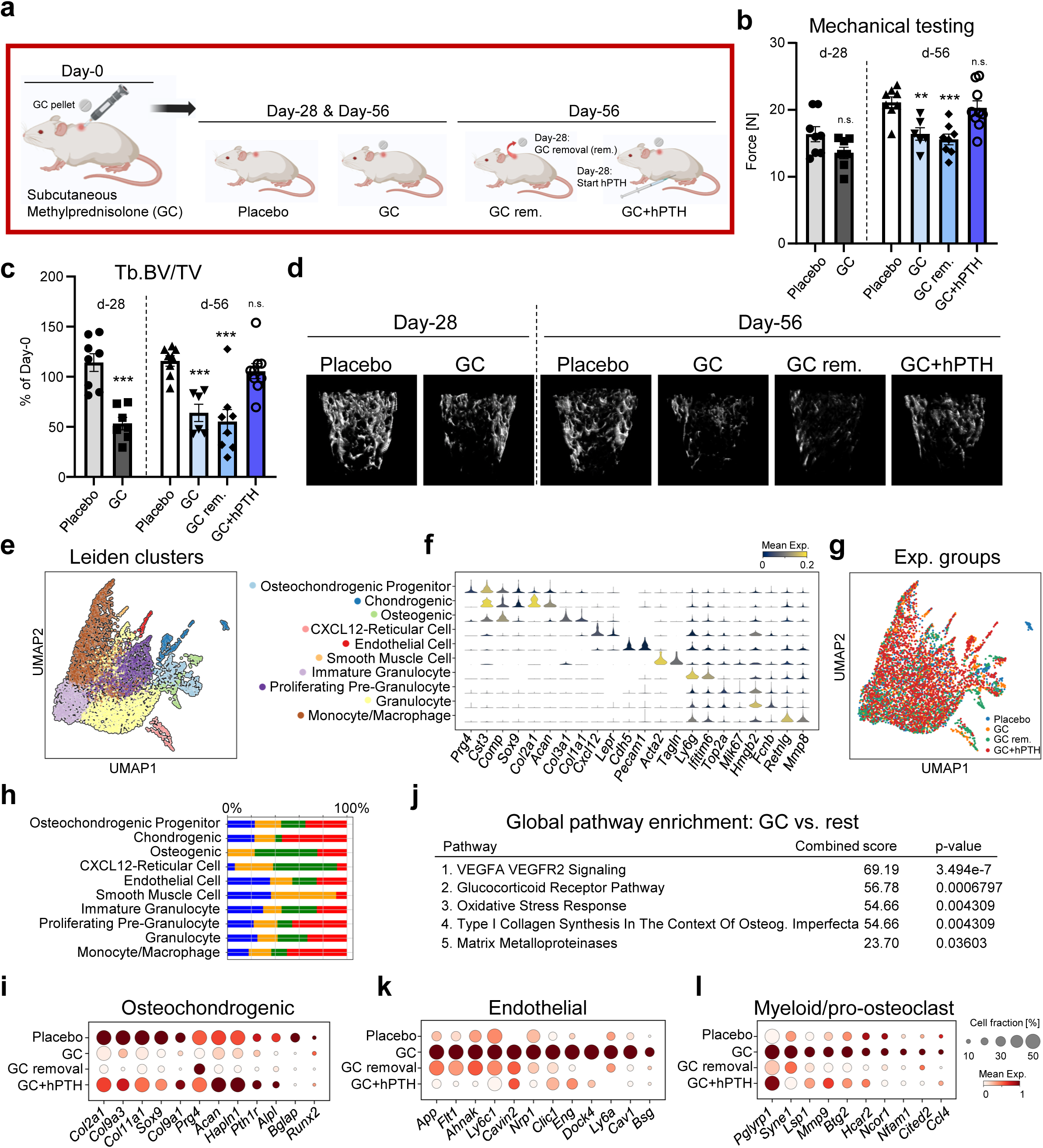
A GC-mediated anti-osteogenic, pro-endothelial, inflammatory shift is reversed by PTH. **(a)** Experimental schematic for exposing mice to continuous glucocorticoids through subcutaneous Methylprednisolone treatment at two timepoints and varying interventions. **(b)** Mechanical strength test of femurs by 3-point bending. **(c)** Quantification of trabecular bone volume per total volume (Tb.BV/TV) shown as percentage change compared to day 0. **(d)** Representative microCT images of distal femur trabecular bone at day-28 and day-56. Data shown as mean ± SEM. Statistical testing between Placebo and other group by unpaired student t-test. *p<0.05, **p<0.01 ***p<0.001, ****p<0.0001. **(e)** UMAP of femur derived single cells from all experimental groups and their distinct clustering by Leiden. **(f)** Specific markers of Leiden clusters determining cellular identity. **(g)** UMAP plot showing cellular clustering labeled by experimental group. **(h)** Bar graphs showing relative abundance of each cell type captured for each experimental group. **(i)** Dotplot showing selected osteochondrogenic gene expression in mesenchymal cell subsets. **(j)** Global pathway enrichment analysis of top 200 expressed genes in mesenchymal cells of GC group using EnrichR. **(k)** Dotplots of endothelial, and **(l)** myeloid gene expression in vascular and hematopoietic cell types, respectively, between different experimental groups.

### GC-exposure drives distinct cellular and molecular changes in bones

Having established our model of GC-induced bone loss, we next sought to derive a more detailed view of the cellular changes of bone tissue. We conducted 10X Chromium single cell RNA-sequencing (scRNAseq) of dissected femurs from the five experimental groups on day 56. Unbiased Leiden clustering analysis of stringently quality filtered single cells established the cellular composition of captured cells for each experimental group (Fig.1e-f). This approach captured the broad heterogeneous composition of the bone/bone marrow (BM) composition including hematopoietic, mesenchymal and endothelial cell types. When we investigated specific differences in the cellular make up of each group, we observed that GCs increased stromal cell populations in the BM including CXCL12 expressing reticular (CAR) cells and smooth muscle cell types (Fig.1g-h). There were also slight changes to committed bone forming and smooth muscle cell types, indicating potential alterations to mesenchymal lineage allocation commitment. Specifically, expression of genes associated with osteogenesis and chondrogenesis were reduced upon GC exposure but showed strong improvement to placebo levels if mice were simultaneously treated with hPTH. Conducting global pathway enrichment analysis with the top 200 differentially expressed genes for each group, revealed that GC exposure led to increases in angiogenesis-related signaling in the BM environment (Fig.1j). Specific gene expression patterns of endothelial genes showed that GC removal and hPTH treatment normalized the expression to placebo levels (Fig.1k).Similarly, while the immune cell compartment composition as not strongly altered by GC exposure, we observed an increase in pro-inflammatory and potentially pro-osteoclastic signaling that was partially reversed by GC removal and hPTH treatment (Fig.1l). In sum, scRNAseq of the bone tissue from the different experimental groups revealed distinct cellular and molecular changes of the BM compartment supportive of alterations at the tissue level. Strikingly, the strongest changes were associated with alterations in angiogenic signaling.

### GCs alter skeletal stem and progenitor function and blood vessel characteristics

Since we observed bone loss and reduced expression of osteochondrogenic genes in mice exposed to GC, we wondered if skeletal stem and progenitor activity might be altered. Therefore, we analyzed the frequency of phenotypic SSCs (CD45^-^Ter119^-^Tie2^-^CD90^-^6c3^-^ CD105^-^CD51^+^) and the directly downstream transient multipotent bone-cartilage-stromal progenitors (BCSPs; CD45^-^Ter119^-^Tie2^-^CD90^-^6c3^-^CD105^+^CD51^+^) using flow cytometry^10^. Results showed that GCs initially (day 28) decreased SSC and BCSP numbers, while extended exposure (day 56) led to an increase in SSCs (Fig.2a & Extended Data Fig.3a-b). However, GC removal led to a significant accumulation of SSCs and BCSP compared to placebo controls. Again, only GC with concomitant hPTH normalized skeletal progenitor abundance to control levels. To further assess functional properties of SSCs, we freshly purified them from bone tissue of each experimental group on day 56 and seeded them for standard in vitro osteogenic and chondrogenic assays. In alignment with micro-CT and transcriptomic data, GCs significantly reduced osteogenic and chondrogenic potential of skeletal progenitors compared to placebo controls which was also not altered upon GC removal assessed after 28 days (Fig.2b). SSCs from bones of mice exposed to GC but treated with hPTH injections showed strong osteochondrogenic activity. These results indicate that skeletal stem and progenitor function is negatively affected by GCs, directly associating it to negative structural and mechanical changes observed for bones.

**Figure 2.**
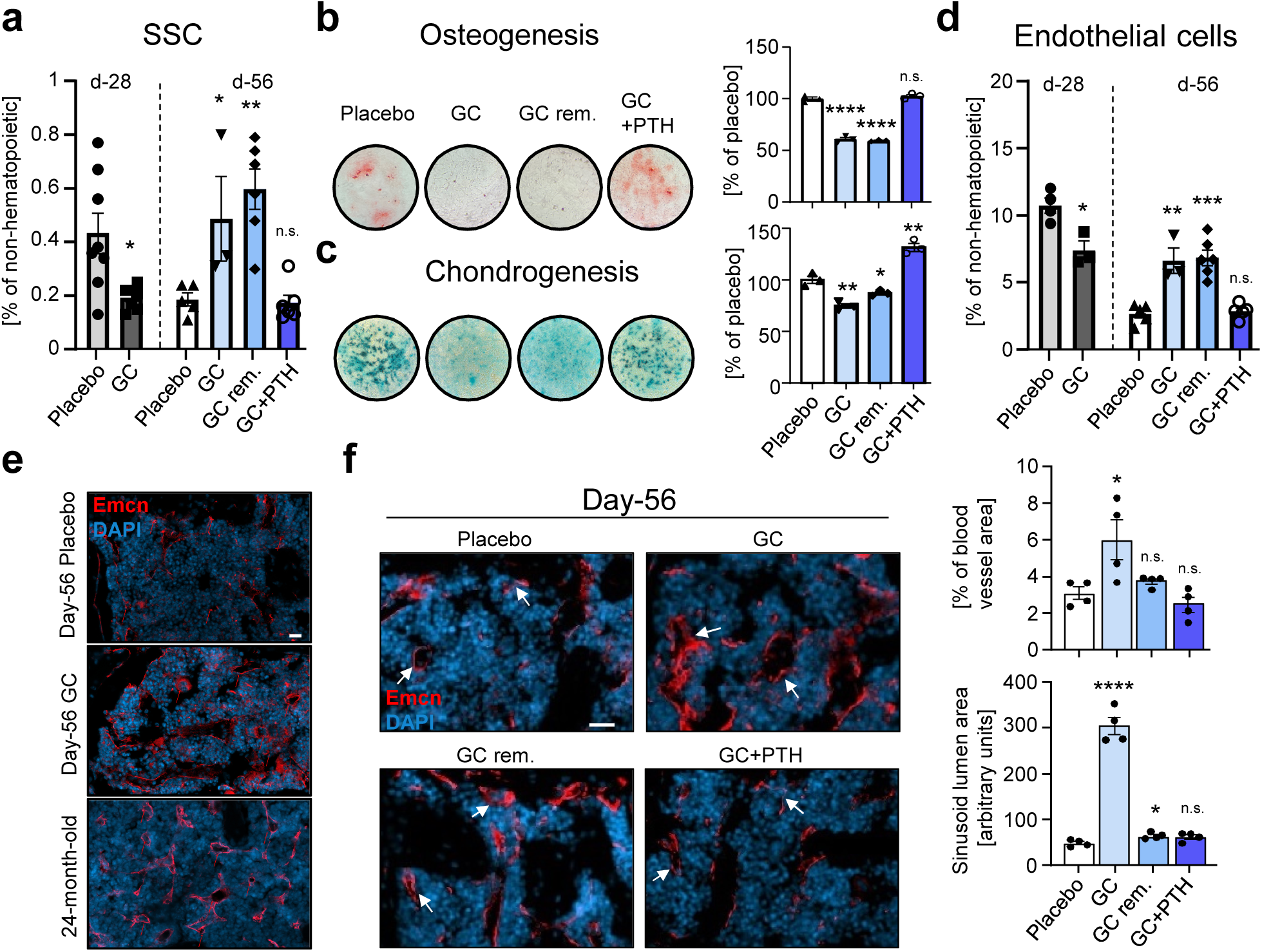
GCs drive impaired skeletal stem and progenitor function and altered bone marrow blood vessel characteristics. **(a)** Flow cytometric based quantification of skeletal stem cell (SSC, CD45-Ter119-Tie2-CD90-6c3-CD105-CD51+) in femurs of experimental groups. n=3-8. **(b)** In vitro osteogenesis (Alizarin Red S stain) and **(c)** chondrogenesis (Alcian Blue stain) assays on purified, primary SSCs. Spectrophotometric quantification of staining (right). n=3. **(d)** Flow cytometric based quantification of endothelial cell populations (CD31+) in femurs of experimental groups. n=3-6 mice per group. **(e)** Immunohistochemistry staining for Endomucin (Emcn) in bone marrow of day-56 placebo and GC treated mice as well as in 24-month-old wild type mice. Arrow heads: sinusoids. **(f)** Representative immunohistochemistry staining for Endomucin (Emcn) in bone marrow of day-56 experimental groups and quantification of blood vessel area (right top) and size of sinusoid lumen (right bottom) based on immunohistochemistry staining for Endomucin. n=4. All data shown as mean ± SEM. Statistical testing between Placebo and other group by unpaired student t-test. *p<0.05, **p<0.01 ***p<0.001, ****p<0.0001. Scale bars, 20µm.

**Figure 3.**
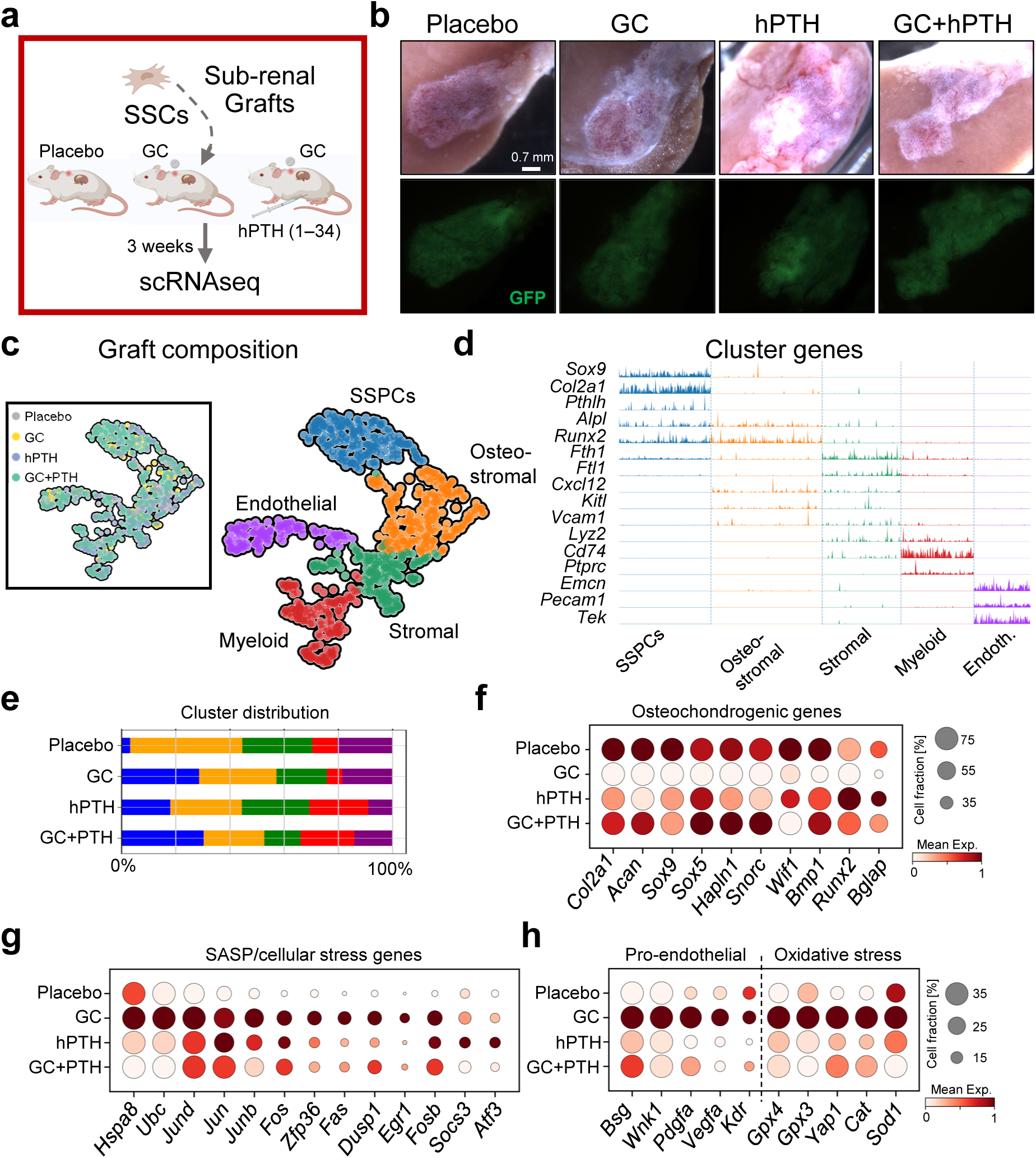
GC alters de novo in vivo bone formation by SSCs and the niches they generate. **(a)** Experimental schematic of renal capsule transplants of freshly purified wild type, GFP-labeled SSCs in mice exposed to placebo, GC or GC+human PTH (hPTH 1-34). **(b)** Light microscopic images (top) and GFP signal of grafts formed beneath the renal capsule of transplanted mice. **(c)** Single cell transcriptomic analysis of formed tissue grafts displayed as UMAP of single cells clustered by Leiden for cell type association and by group (top left). SSPC: Skeletal stem and progenitor cells. **(d)** Tracksplot of specific markers of Leiden clusters determining cellular identity. **(e)** Bar graphs showing relative abundance of each cell type captured for each experimental group. **(f)** Dotplot showing selected osteogchondrogenic (left) genes in SSPC cluster between different experimental groups. **(g)** Dotplot showing SASP (senescence associated secretory phenotype)/cellular stress related gene expression in all cells between different experimental groups. **(h)** Dotplot showing pro-endothelial and oxidative stress related gene expression between different experimental groups.

Based on scRNAseq results, we also wondered how endothelial numbers and composition maybe altered. Flow cytometric analysis revealed a similar pattern as seen with SSCs. While after 28 days, endothelial cell numbers were reduced with GCs compared to the placebo group, longer GC exposure drove an accumulation of endothelial cell presence that was only normalized to placebo levels when mice received hPTH with GCs (Fig.2d & Extended Data Fig.3c). When we stained histological sections of femur bones after 56 days of GC exposure with the endothelial marker Endomucin, we observed an increase of blood vessel abundance as well as distinct morphological changes that resembled the endothelial compartment of aged (24-month-old) mice (Fig.2e). Previous work has reported that inflammatory stress to the BM environment is connected to increased numbers of sinusoids with dilated lumina^11^. Quantitative analyses showed that at day 56 of GC exposure there were significantly more Endomucin-positive sinusoidal blood vessels with increased lumen area in the BM compared to placebo controls (Fig.2f & Extended Data Fig.3d). The blood vessel area and dilation of lumens remained at placebo levels when mice were concomitantly treated with hPTH. In summary, GCs mediate specific changes to bone-forming lineage cells and angiogenic processes that can be reversed by co-stimulation with daily hPTH, suggesting a direct connection between SSCs and endothelial cells.

### GCs alter SSC-mediated ossicle formation

To determine if GC-mediated alterations were connected through changes observed in SSC and endothelial cell activity, we assessed SSC-specific bone formation characteristics via renal capsule transplantation^12^. To that end, we freshly FACS purified equal numbers of SSCs from ubiquitous GFP-reporter mice and transplanted them beneath the renal capsule of wild type C57BL/6 mice. We randomly assigned mice to four groups, i.e., a control group receiving placebo pellets at the time of surgery, a group with hPTH treatments but no GCs as well as two groups receiving GC release pellets. One of the GC groups received daily hPTH over the 21 days of the experiment (Fig.3a). At that time, we isolated tissue grafts and processed them for scRNAseq. Generated ossicles of all groups contained mesenchymal, endothelial and hematopoietic cell types (Fig.3b-e). Mice treated with GCs alone presented with a higher number of undifferentiated skeletal stem and progenitor cells (SSPCs) than placebo treated mice. The expression of osteogenic and chondrogenic gene programs was also decreased with GC treatment alone but was rescued by hPTH treatment (Fig.3f). Analyzing gene expression patterns of all cells, GC exposure led to high expression of genes related to cellular stress including those associated with the senescence-associated secretory phenotype (SASP) and oxidative stress, supporting the idea that GCs appear to drive an aging-like and pro-inflammatory microenvironment) (Fig.3g). Interestingly, genes involved with blood vessel recruitment and formation were highly increased in the presence of GCs which also was associated with increased oxidative stress gene expression (Fig.3h). Once again, hPTH added to the GC treatment was able to reverse gene expression patterns to placebo control levels. Since SSC derived ossicle formation strongly depends on host endothelial blood vessel recruitment, these results support a co-regulation of SSC and endothelial lineages during GC exposures and provided a list of molecular interaction partners.

### GC-induced SSC derived Basigin impairs bone forming and vascular network properties

Among the multiple pro-endothelial genes upregulated upon GC exposure (*Kdr/Vegfr*, *Vegfa*, *Pdgfa*, *Wnk1*) we identified *Basigin* (*Bsg*) to be highly expressed in accumulated SSPCs, and was reversed by hPTH treatment (Fig.1k, 3h & 4a). We hypothesized that Basigin, a known activator of smooth muscle cell proliferation, might in part drive GC-mediated aberrations in the BM environment. To further explore this concept, we conducted functional tests in primary human SSCs (hSSCs) and in the human VeraVec HUVEC endothelial cell line. We collected 48h supernatant from hSSCs either lentivirally overexpressing Basigin or vector controlled to directly tie SSPC derived Basigin expression to changes in endothelial cell activity. Then we assessed the effect of the supernatants of both groups during in vitro tube formation and wound scratch assays of VeraVec endothelial cells. Strikingly, in the presence of high levels of Basigin in the supernatants, blood vessel architecture was significantly impaired, as indicated by reduced mesh, tube and node numbers compared to the control (Fig.4b-c). We also found that Basigin impaired endothelial cell migration in a wound scratch assay (Fig.4d). In addition, elevated Basigin levels increased ROS generation in cultured endothelial cells (Fig.4e), confirming scRNAseq readouts. Furthermore, overexpression of Basigin in hSSCs drove increased colony forming ability, is a measure of proliferative activity, while it impaired in vitro osteogenic and chondrogenic differentiation compared to controls (Fig.4f-h). Finally, we subcutaneously transplanted equal numbers of hSSCs either overexpressing Basigin or control into immunodeficient NOD scid gamma (NSG) mice to determine if ossicles formed displayed differences in osteogenesis and angiogenesis in vivo (Fig.4i-j). In line with a recent report on the effect of GCs on fracture healing^13^, we observed reduced bone remodeling dynamics in grafts derived from Basigin-overexpressing cells that were less mineralized and showed low bone resorption activity by host-derived osteoclasts (Fig.4k-l). Interestingly, the shortened morphology of recruited blood vessels of the Basigin-SSC grafts mirrored the GC-induced phenotype observed in femurs (Fig.4m). These results establish a direct connection between SSC-derived Basigin and detrimental effects of skeletogenesis and blood vessel architecture caused by GCs that is transferable to human cells.

**Figure 4.**
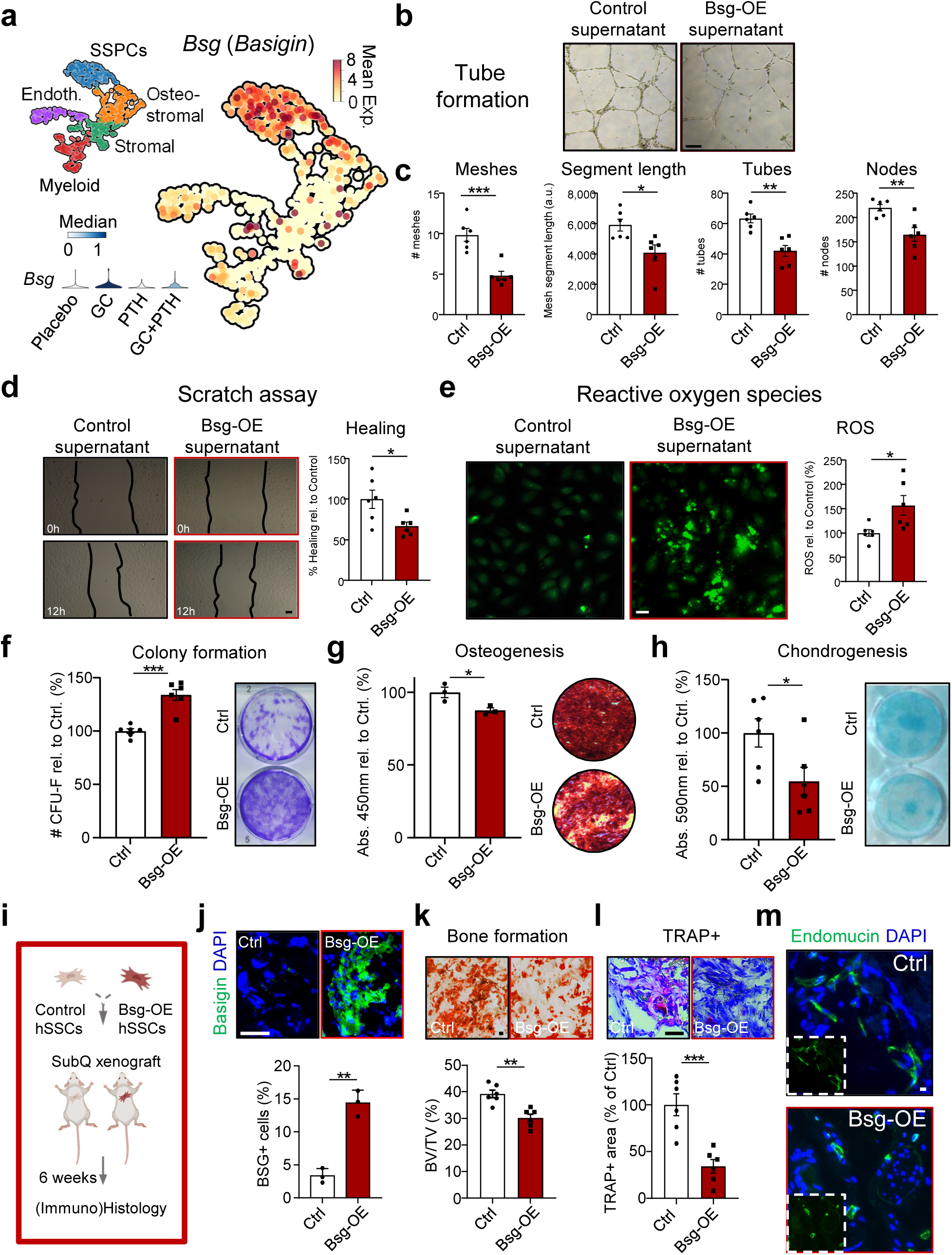
Basigin overexpression alters human skeletal lineage dynamics that impair endothelial function. **(a)** Single cell transcriptomic analysis of tissue grafts from mouse SSCs either exposed to placebo, GC, human PTH or GC+human PTH (hPTH 1-34) displayed as UMAP showing high expression of Basigin in SSPCs. SSPC: Skeletal stem and progenitor cells. Left bottom: Violin plot showing increased expression of *Bsg* in GC group across all cells. **(b**) Representative image of tube formation assay by VeraVec endothelial cells exposed to control supernatant or supernatant of human SSCs overexpressing Basigin. **(c)** Quantification of ImageJ-based tube formation analysis. n=6. **(d)** Representative brightfield images of endothelial scratch assay (left) and its quantification (right). n=6. **(e)** Measurement of reactive oxygen species in cultured endothelial cells after 24h supernatant exposure. Left: representative fluorescence images. Right: Quantification of fluorescence signal. n=6. All experiments with supernatant from at least two donors and two independent experiments. **(f)** Colony forming unit ability of primary human SSCs either overexpressing control vector or Basigin. CFU-Fs stained by Crystal violet. n=6 **(g**) In vitro osteogenesis (Alizarin Red S stain) by hSSCs of the same groups. n=3. **(h)** In vitro chondrogenesis (Alcian Blue stain) assays of the same groups. n=6. **(i)** Schematic of subcutaneous transplant approach. **(j)** Immunohistochemistry of Basigin (green) expression in SSC-generated grafts with corresponding quantification. n=3. **(k)** Representative Alizarin Red S staining of sectioned grafts and quantification of mineralized tissue in grafts containing transplanted human SSCs as assessed by Alizarin Red S staining. n=6. **(l)** Representative TRAP staining of sectioned grafts and quantification of TRAP-positive area of grafts. n=6. **(m)** Representative immunohistochemistry staining for Endomucin and DAPI of sectioned graft. Small insert shows Endomucin staining without DAPI. All data shown as mean ± SEM. Statistical testing between Placebo and other group by unpaired student t-test. *p<0.05, **p<0.01 ***p<0.001, ****p<0.0001. Scale bars, 50 µm

### Antibody-mediated blockade of hSSC-derived Basigin reverses impaired endothelial function and bone formation in vitro

Next, we investigated whether the negative effects of Basigin-overexpression in human SSCs could be pharmacologically reversed. To that end, we treated human SSCs with a monoclonal antibody against Basigin (aBSG) and collected supernatant to test its effect on vascular modeling. Indeed, the detrimental paracrine effect on human endothelial tube formation in the presence of supernatant from Basigin-overexpressing SSCs was mostly returned to control levels with antibody treatment (Fig.5a). Similarly, VeraVec cells exposed to Basigin and treated with aBSG performed similarly to controls in wound scratch assays (Fig.5b). Strikingly, the Basigin induced osteogenesis-impairing effects due to overexpression in SSCs were rescued when we either exposed the cells to hPTH or aBSG (Fig.5c), supporting Basigin antibody blockade as a novel strategy to prevent or reverse GC-induced SSC dysfunction mediated bone loss.

**Figure 5.**
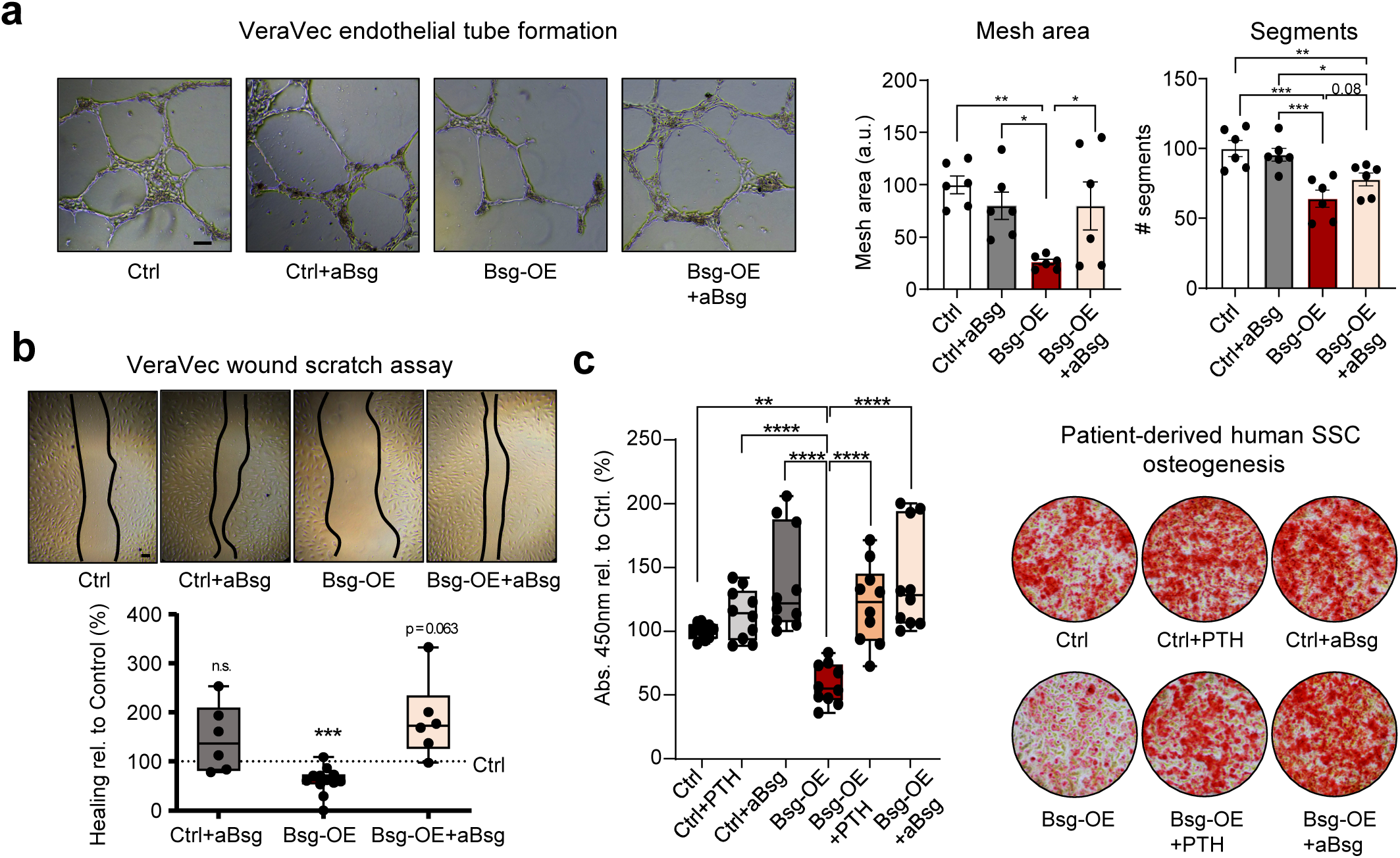
Antibody blockade of Basigin rescues GC-induced endothelial and skeletal impairments in vitro. **(a)** Tube formation assay of human VeraVec endothelial cells treated with supernatant from different experimental groups. **(b)** Representative brightfield images of endothelial scratch assay (top) and its quantification displayed as boxplots (bottom) after 12h.Endothelial cells were treated with supernatant of cultured hSSCs treated as shown. n=6-12. **(c)** In vitro osteogenesis of patient-derived human SSCs. Left: Boxplots showing quantification of Alizarin Red S staining. Right: Representative images of Alizarin Red S staining. Ctrl: control media only; Ctrl+PTH: control media with PTH 1-34 treatment 6h before media change; Ctrl+aBsg: control media with Basigin antibody treatment. Bsg-OE: Lentivirally Basigin overexpressing human SSCs with control media. Results generated using cells from at least two independent donors in at least two independent experiments. All data shown as mean ± SEM. Statistical testing in a,c by one-way ANOVA with Fisher-LSD test, in b by Wilcoxon signed-rank test. *p<0.05, **p<0.01 ***p<0.001, ****p<0.0001. Scale bars, 100 µm

### Blocking Basigin overexpression by antibody therapy in vivo rescues detrimental bone loss induced by GCs

To confirm our in vitro findings, we next tested whether antibody blockade of Basigin can prevent GC-induced bone loss in mice. We compared placebo treated control mice with mice that were exposed to GCs for 28 days. Of these GC-treated mice we looked at mice receiving no additional therapy or treated with hPTH or aBSG throughout that time period (Fig.6a). Histological analyses of femoral bone sections at day 28 showed that hPTH and aBSG treatments reduced Basigin expression seen in the GC only group (Fig.6b). Also, the reduction in Basigin levels in these treatment groups correlated with trabecular bone volume, osteoclast numbers and bone marrow endothelial morphology of placebo controls (Fig.6c-e), prevented the GC-induced short-term reduction in SSC frequency (Fig.2a) and reinstated their in vitro osteogenic potential (Fig.6f-g). While we observed a GC-driven shift towards an increase in circulating myeloid cell types in blood which was reversed by aBSG treatment, we did not detect any significant differences on the hematopoietic stem and progenitor compartment in the bone marrow (Fig.6h-I & Extended Data Fig.4). Thus, administration of aBSG during GC treatment prevents skeletal maladaptation.

**Figure 6.**
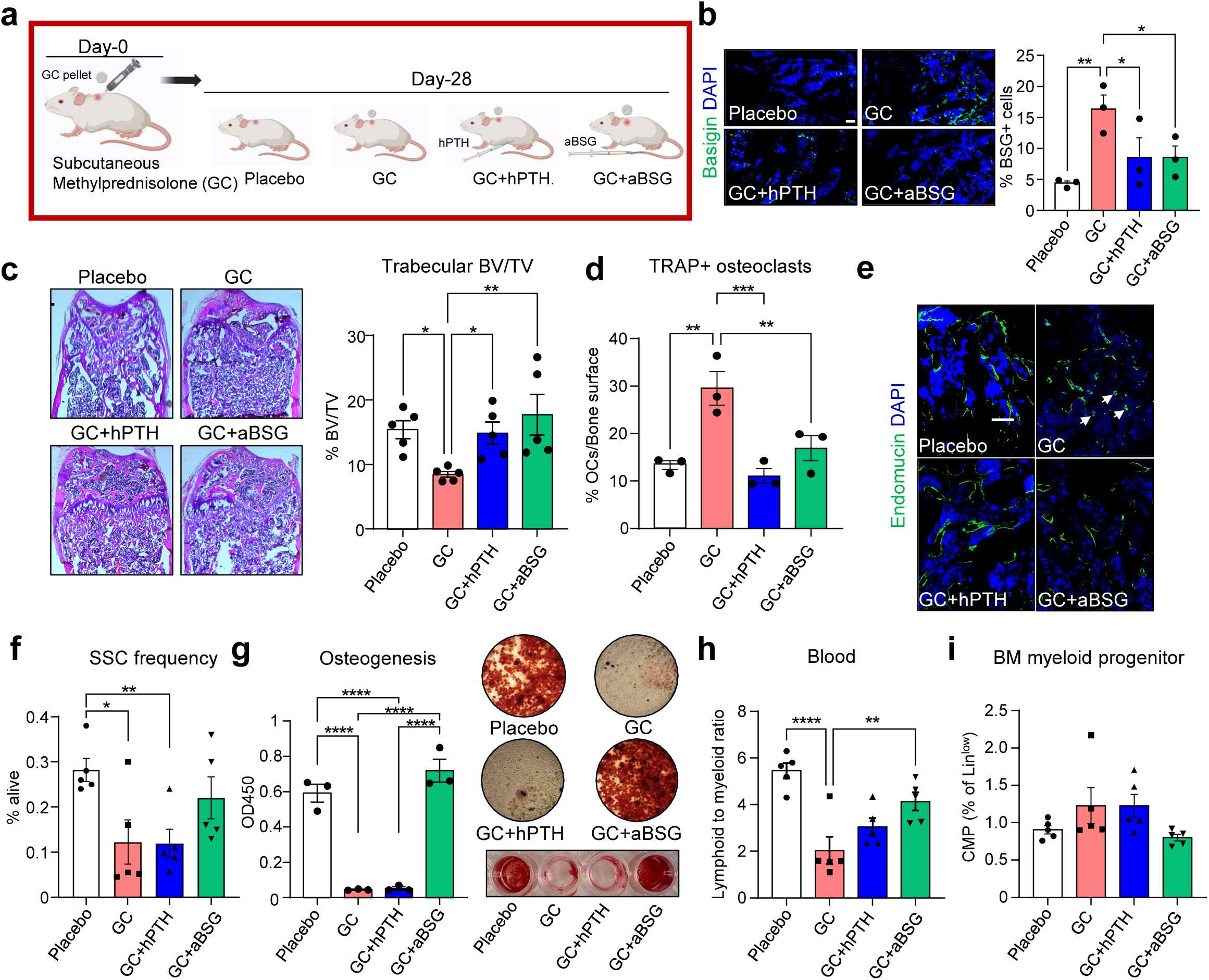
Basigin antibody blockade reverses GC-induced skeletal changes. **(a)** Schematic of experimental setup. **(b)** Representative immunohistochemistry images and quantification of Basigin expression in the different experimental groups. **(c)** Representative H&E stain of distal femurs of experimental groups with quantification of trabecular bone area below the growth plate region. n=5. **(d)** TRAP staining and quantification of femur bones from all experimental groups. **(e)** Representative images of Endomucin-positive endothelium in the bone marrow environment. Arrows: small sinusoids. **(f)** Flow cytometric analysis of femur bones of experimental groups for skeletal stem cells (SSCs). **(g)** In vitro osteogenic differentiation of bone marrow stromal cells derived from experimental groups stained with Alizarin Red. **(h)** Flow cytometric analysis of blood from mice of different experimental groups showing lymphoid to myeloid ratio. **(i)** Flow cytometric analysis of bone marrow (BM) common myeloid progenitor cells (CMPs) from mice of different experimental groups. All data shown as mean ± SEM. Statistical testing by one-way ANOVA with Fisher-LSD test. *p<0.05, **p<0.01 ***p<0.001, ****p<0.0001. Scale bars, 50 µm

### Basigin antibody therapy improves bone mass in aged mice independent of sex

Since Basigin has been reported to be a therapeutic target for a number of pro-inflammatory and pro-fibrotic conditions, we asked whether aBSG could also reverse age-related bone loss, i.e., osteoporosis^14,15^. Indeed, immunohistochemistry analysis showed higher expression of Basigin in bone marrow of old mice, in particular in females (Fig.7a). When we treated 2-year-old female and male mice with aBSG three times/week at 1 mg/kg for four weeks we found a significant improvement in trabecular BV/TV of long bones compared to IgG control treated mice (Fig.7b-c). Vertebral L5 bone mineral density at 14 days and BV/TV at 28 days of treatment measured by DEXA and micro-CT, respectively, demonstrated an anabolic effect on the skeleton by aBSG independent of sex (Fig.7c-d). Interestingly, SSCs of the same mice showed higher in vitro osteogenic potential whereas bone marrow culture readouts and histological analysis also suggested an increase in bone resorptive activity with aBSG (Fig.7f-g). This was accompanied by endothelial restoration but no detectable changes in the frequency of skeletal and hematopoietic stem and progenitor cells (Extended Data Fig.5). Collectively, this indicated that aBSG treatment boosts bone remodeling in aged mice in a sex-independent manner leading to a net gain in bone mass through higher bone-forming activity.

**Figure 7.**
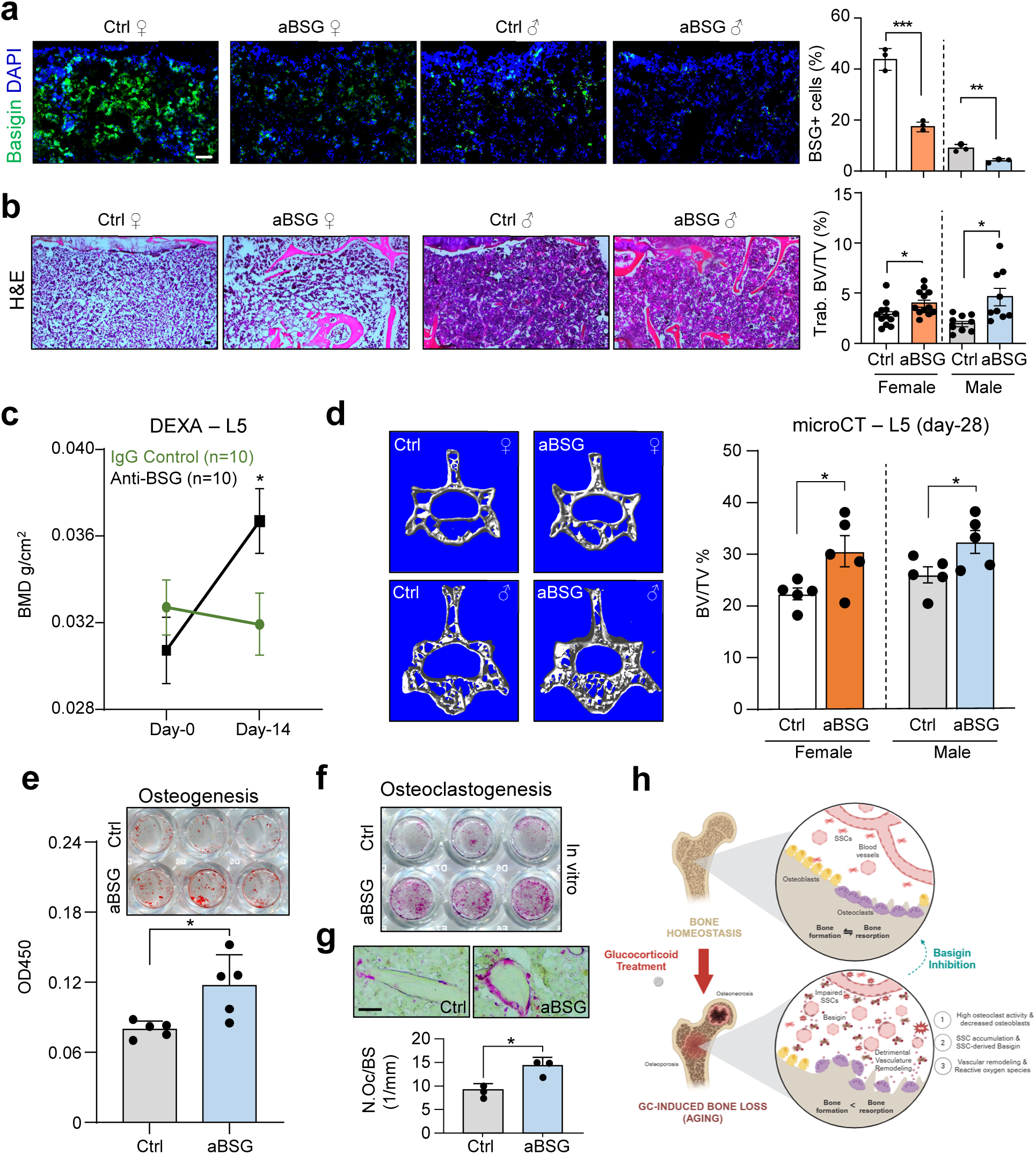
Antibody blockade of Basigin in aging mice improves bone parameters. **(a)** IHC of Basigin expression in tibias of 24-month-old mice. Right: Histology-based quantification Basigin-expression cells per analyzed area. **(b)** H&E staining of proximal tibia regions below the growth plate. Right: Histology-based quantification of tibial trabecular BV/TV. **(c)** DEXA bone mineral density (BMD) measurements of vertebral L5 at treatment starting day-0 and at day-14 of treatment. n=5. **(d)** Micro-CT measurement of vertebral L5 BV/TV at 4-week of treatment. n=5. **(e)** Osteogenic differentiation of SSCs derived from experimental groups at day-28. Top: representative images of well plates with Alizarin Red staining. Bottom: quantification. n=5. **(f)** TRAP staining of BM-derived osteoclasts in vitro. **(g)** TRAP quantification in tibia sections of experimental groups. N.Oc/BS: number of osteoclasts per bone surface. n=3. **(h)** Graphical summary of experimental findings. All data shown as mean ± SEM. Statistical testing between Placebo and other group by unpaired student t-test. *p<0.05, **p<0.01 ***p<0.001, ****p<0.0001. Scale bars, 50 µm

## Discussion

While it has been established that GCs increase osteoclastogenesis and alter osteogenesis and angiogenesis, the mechanism related to vascular changes has not been appreciated. In this study, we identified a previously unknown connection between skeletal stem cell progenitor function and angiogenesis. Continuous treatment with GCs promoted alterations to the BM blood vessel network by direct actions on SSCs that were locked in an undifferentiated state that prevented osteochondrogenic lineage commitment. We uncovered a novel potential mechanism by which Basigin, expressed by SSCs, when exposed to GCs alters endothelial function and remodels BM angiogenesis driving cellular stress and promoting bone loss through reduced formation and increased bone resorption (Fig.7h). Single cell gene expression data suggests that a consequence of enhanced Vegfr signaling may be driving a pathological blood vessel phenotype through emphasized oxidative stress. The observation that this pathological signaling mechanism is reversable presents novel therapeutic vantage points to counter GC-induced bone loss and osteonecrosis. Concomitant administration of hPTH or aBSG prevented bone loss observed with GC exposure. It remains to be determined if this reversal has any impact on the intended clinical effects of GC therapy.

Our examination of gene expression by scRNAseq of the bone tissue from the different experimental groups revealed distinct cellular and molecular changes of the BM compartment. GCs caused a reduction in osteochondrogenic gene expression associated with decreased bone formation, but also an increase in pro-inflammatory myeloid signaling, potentially promoting osteoclastogenesis. This observation is consistent with the bone cell activity changes observed in the GC treated subjects as GC treatment reduces bone formation and increases the number and activity of osteoclasts^8,9^. However, the strongest changes were associated with alterations in angiogenic signaling which may correlate with the association GC induced osteonecrosis (ON). While epidemiologic studies find GCs increase the risk of ON, most commonly at the femoral head, the pathology of the ON lesion is that of reduced blood vessel density and increased adipose tissue within the bone marrow^8^. Our observation of both the increased angiogenic gene expression and tissue-based quantification of increased number and altered size/structure of bone blood vessels in the presence of GCs, led us to further investigate a potential mechanistic connection. We identified Basigin, to be highly expressed in SSPCs during the GC treatment, and its exposure to endothelial cells resulted in irregularities in the connectivity and thickness of the nascent blood vessels. This observation may be relevant to GC-induced ON, as the altered morphology of these nascent blood vessels may reduce the ability to deliver hemoglobin/oxygen to the bones. In certain areas of the skeleton like the femoral head, this may lead to increased cell death. Additional studies are needed to confirm this observation.

Interestingly, after 28 days of GC treatment, the addition of hPTH resulted in a nearly complete reversal of the GC effects to the baseline levels including osteogenesis, angiogenesis, and Basigin gene expression. These results complement clinical studies reporting that the addition of hPTH to subjects with GC-induced osteoporosis leads to an increase in trabecular bone mass and biochemical markers of increased bone formation. Furthermore, meta-analyses of multiple studies have reported improved bone strength^6,16^. However, the effect of hPTH on angiogenesis in the presence of GCs has not been investigated. Our study demonstrates that GC and hPTH together reduced Basigin expression, restored blood vessel number and morphology, as well as osteochondrogenic differentiation of SSCs. While hPTH has previously been shown to be osteoanabolic and to direct blood vessels toward the bone forming surface, it has not been found to be pro-angiogenic^17^. Even though GCs and hPTH likely act on multiple skeletal lineage cell populations, including osteoblasts, our results provide evidence for their crucial role in regulating SSC activity. Recognizing that many phenotypic changes in response to GC treatment resembled age-associated skeletal changes, GC-induced bone loss might underlie overlapping mechanistic processes^18^. Our studies with aBSG support that notion. Future work will have to inquire if aBSG might be a superior therapeutic approach to hPTH to prevent GC-mediated bone loss, given the limited amount of time hPTH can be administered for beneficial clinical outcomes.

We observed that the withdrawal of GCs on day 28 did not restore bone mass to baseline levels by day 56. Elevated GCs result in a reduction in the release of GCs from the hypothalamus^19^, and our 28-day GC treatment may have sufficiently suppressed the hypothalamic-pituitary-adrenal (HPA) axis such that the 28-day treatment period was insufficient to see recovery of the skeletal tissues. In support of this observation, both osteogenesis and angiogenesis measurements for this group were similar to those of the GC only group at 56 days. More studies will need to investigate the HPA axis recovery in this treatment group to contextualize our current findings.

Finally, we made the novel discovery that Basigin from SSPCs might be partially responsible for maladaptive coupling between skeletal and angiogenic cell lineages of bone. Related to that, Basigin has been considered an innovative marker in the context of cancer stem cells^20^, but little is known about Basigin in the realm of skeletal biology. Two studies report that high levels of Basigin promote osteoclastogenesis through NFATc1 signaling^21^ and its involvement in alveolar bone remodeling and soft tissue degradation^22^. It also acts as a potent stimulator of Interleukin-6 secretion in multiple cell lines that include monocytes^23^, altogether supporting our observation of increased inflammatory signaling and bone resorption activity in GC treated animals with high SSPC specific Basigin expression. Fitting with the observation of SSC accumulation and altered cytokine expression, Basigin has also been found to contribute to tumor progression by stimulation of proliferation and elevated growth factor secretion as well as liberation through its matrix metalloprotease activity^24^. Lastly, Basigin’s role as a coreceptor for vascular endothelial growth factor receptor 2 (Kdr/Vegfr2) in endothelial cells enhancing its Vegfra-mediated activation and downstream signaling, clearly supports our findings connecting it to blood vessel formation ^25^. Intriguingly, we could show that antibody treatment against Basigin mitigates both its detrimental autocrine and paracrine effects on skeletogenesis and vascular modeling, respectively. Excitingly, results in aged mice suggest a potential for aBSG as an anti-osteoporotic drug. In summary, our report is the first to report the detrimental role of Basigin signaling in osteoangiogenic coupling of bones and by provides a new therapeutic target to potentially prevent or reverse skeletal health decline. Future studies will determine how modifying Basigin in vivo can prevent both bone loss and osteonecrosis in the presence of GCs.

### Materials & Methods

#### Animals

All mouse experiments complied with all relevant ethical regulations and were conducted under approved protocols by the Institutional Animal Care and Use Committee of Stanford University and the University of California Davis. Mice were maintained at animal facilities in accordance with institutional guidelines. Mice were given food and water ad libitum and housed in temperature-, moisture- and light-controlled (12-h light-dark cycle) micro-insulators. Unless otherwise specified, all experiments were conducted using 3-month-old Balb/cJ male mice, purchased from Jackson Laboratories (Strain#:000651). To study the effects of glucocorticoids on bone biology, Methylprednisolone 5 mg 60-day release pellets or vehicle (cat#SG-241, Innovative Research of America, Sarasota, FL, USA) were implanted under the skin near the lumbar spine at day 0, and pellets were maintained for 56 days or removed after 28 days as indicated. Human parathyroid hormone 1-34 (hPTH 1-34) was purchased from Sigma (cat#:P3796), and the treatment was administered subcutaneously at 40 μg/kg, 5x/week for four consecutive weeks. For 28-day studies with concomitant hPTH (40 μg/kg/day, subcutaneously 5x/week) or aBSG (1mg/kg, i.v. 3x/week) treatment mice were implanted with Methylprednisolone 2.5 mg 21-day release pellets or vehicle control pellets. Aged (24-months) male and female C57BL/6 from were ordered from NIA and treated i.v. with aBSG or IgG controls (1mg/kg, 3x/week for four weeks). For renal capsule transplantation experiments cells from three-month old male GFP reporter mice (C57BL/6-Tg(CAG-EGFP)1Osb/J; JAX: 003291) were transplanted into male B6 mice (C57BL/Ka-Thy1.1-CD45.1; JAX: 000406). At the same day mice were transplanted with GC pellets (2.5 mg 21-day release pellets or vehicle) and hPTH treatment was initiated (40 μg/kg/day, 5x/week for 3 weeks). Three-month old immunodeficient NSG mice (NOD.Cg-Prkdcscid Il2rgtm1Wjl/SzJ; JAX: 005557) were used for human cell transplantation studies.

#### Human primary cells and cell lines

Human skeletal stem cells (hSSCs) were obtained from fracture callus (day 3-5 after injury) tissues during open reduction and internal fixation procedures. Procurement and handling were in accordance with the guidelines set by the Stanford University Institutional Review Board (IRB-35711) and the UC Davis Institutional Review Board (IRB-1997852). Informed consent was not required as samples were considered biological waste. No restrictions were made regarding the race, gender, or age of the specimen’s donor. Following excision, all specimens were placed on ice, and hSSCs were isolated as described below. Early passage (3-5) VeraVec HUVEC cells were used for endothelial assays as further explained below.

#### Micro-computed tomography and dual-energy x-ray absorptiometry (DEXA) analysis

Soft tissue-free femurs were scanned within 2 h of dissection using a Bruker Skyscan 1276 (Bruker Preclinical Imaging) with a source voltage of 85 kV, a source current of 200 μA, a filter setting of AI 1 mm, and pixel size of 17.5μm at 2016 × 1344. Reconstructed samples were analyzed using CT Analyser (CTan) v.1.17.7.2 and CTvox v.3.3.0 software (Bruker). Anatomical landmarks as per ASBMR guidelines^26^ were used to set region of interest for analyzing trabecular and (200 consecutive sections) cortical (100 consecutive sections) bone parameters. Subsequent mechanical strength testing was conducted using 3-point bending. The maximum load (N) sustained prior to fracture was recorded ^27^. Whole-body DEXA imaging was performed at baseline (day of surgery) and 14 days post-surgery to determine lumbar spine (L5) BMD. Mice were anesthetized with isoflurane and placed in a cabinet x-ray system (Mozart®, Kubtec Medical Imaging) for analysis.

#### Histology

Soft-tissue-free specimens were fixed in 4% PFA at 4 °C overnight. Samples were decalcified in 400 mM EDTA (EDTA) in PBS (pH 7.2) at 4 °C for 2 weeks with a change of EDTA every other day. The specimens were then dehydrated in 30% sucrose at 4 °C overnight. Specimens were embedded in optimal cutting temperature compound (OCT) and sectioned at 5 µm. Representative sections were stained with freshly prepared hematoxylin and eosin (H&E) or Tartrate-resistant acid phosphatase (TRAP). Immunofluorescence on sections of cryopreserved long bone and ectopic bone specimens were incubated with 3% Bovine Serum Albumin in Tris Buffered Saline (TBS) for 1h. Then, samples were probed with primary antibody (Endomucin; 1:200; cat# sc-65495, Santa Cruz) diluted in 1% BSA/PBS and incubated in a humidified chamber at 4°C overnight. The specimens were washed with PBS three times. Secondary antibody was applied for 15 minutes at room temperature in the dark. Specimens were also incubated with 1 µg/ml of DAPI for 10 mins and then washed twice. The specimens were then mounted with a coverslip using Fluoromount-G and imaged. ImageJ (http://wsr.imagej.net/distros/osx/ij152-osx-java8.zip; RRID:SCR_003070) was used to quantify Endomucin-positive blood vessel/sinusoid lumen area. H&E stains were analyzed using image deconvolution according to Landini et al.^28^. For quantification, the software ImageJ (National Institutes of Health, Bethesda, MD, USA) was used on images of H&E sections. The bone marrow area spanning 1 mm below the growth plate between cortical bone region was selected and the image deconvolution plugin, Colour Deconvolution 2, was ran on the selected region. The H&E 2 filter within the plug-in was applied. The color threshold on panel “Colour_2” was adjusted to represent the Eosin-stained bone regions. To measure the BV/TV, the partial (bone) area and total area and were analyzed.

#### Bone Histomorphometry Analysis

Mice were injected with 20 mg/kg of Calcein (Sigma-Aldrich, St. Louis, MO, USA) 8 and 2 days before euthanasia. Histological sections of bones were prepared as described above. A standard sampling site in the secondary spongiosa of the distal metaphysis was established. Mineral apposition rate (MAR), bone formation rate (BFR) and osteoclast number/bone surface (Oc/BS) were calculated.

#### Flow cytometric isolation of skeletal progenitor cells

SSC lineage populations were isolated using cell-surface-marker profiles as previously described ^10,12^. In brief, femurs were dissected, cleaned of soft tissue and crushed using mortar and pestle. Then, the tissue was digested in M199 (cat#11150067, Thermo Fisher Scientific) with 2.2 mg/ml collagenase II buffer (cat#C6885, Sigma-Aldrich) at 37 °C for 60 min. Dissociated cells were strained through a 100-μm nylon filter, washed in staining medium (10% fetal bovine serum (FBS) in PBS) and pelleted at 200*g* at 4°C. The cell pellet was resuspended in staining medium and red blood cells were depleted via ACK lysis for 5 min. The cells were washed again in staining medium and pelleted at 200*g* at 4°C. Then, the cells were prepared for flow cytometry with fluorochrome-conjugated antibodies. For mouse SSC lineages: CD90.1 (Thermo Fisher, 47–0900), CD90.2 (Thermo Fisher, 47–0902), CD105 (Thermo Fisher, 13– 1051), CD51 (BD Biosciences, 551187), CD45 (BioLegend, 103110), Ter119 (Thermo Fisher, 15–5921), Tie2 (Thermo Fisher, 14–5987), 6C3 (BioLegend, 108312), streptavidin PE-Cy7 (Thermo Fisher, 25–4317), Sca-1 (Thermo Fisher, 56-5981), CD45 (Thermo Fisher, 11–0451), CD31 (Thermo Fisher, 12-0311) and CD24 (Thermo Fisher, 47–0242). For human SSC lineages: CD45 (BioLegend, 304029), CD235a (BioLegend, 306612), CD31 (Thermo Fisher Scientific, 13-0319), CD202b (TIE-2) (BioLegend, 334204), streptavidin APC-AlexaFlour750 (Thermo Fisher, SA1027), CD146 (BioLegend, 342010), PDPN (Thermo Fisher Scientific, 17-9381), CD164 (BioLegend, 324808) and CD73 (BioLegend, 344016). Flow cytometry was conducted on a FACS Aria II Instrument (BD BioSciences) using a 70-μm nozzle in the Shared FACS Facility in the Lokey Stem Cell Institute (Stanford). The skeletal stem-cell lineage gating strategy was determined using appropriate isotype and fluorescence-minus-one controls. Propidium iodide staining was used to determine cell viability. Cells were sorted for purity. Flow cytometric analysis was conducted using FlowJo (FLOWJ LLC, v10.10). Blood panels and bone marrow analysis was conducted on a CYTEK Aurora Sorter. Blood was stained with Ter119-PE-Cy5 (116210, BioLegend), CD45-FITC (11-0451, Invitrogen), B220-APC-Cy7 (103224, BioLegend), CD11b-PE-Cy7 (101216, BioLegend), CD3-APC (100236, BioLegend), Gr1-BV711 (108443, BioLegend). CD11b^+^ and Gr1^+^ cells within CD45+ were considered myeloid lineage, while CD45+CD11b-Gr-CD3+ and B220+ were marked as lymphoid lineage. HSC lineage populations were stained with Lineage cocktail-Pacific Blue (133310, BioLegend), CD127-BV711 (135035, BioLegend), CD117-APC-Cy7 (105826, BioLegend), Sca1-APC (160904, BioLegend), CD16/32-PE (156606, BioLegend), CD34-BV786 (742971, BD Biosciences), CD135-PE-Cy5 (135312, BioLegend), CD150-BV510 (115929, BioLegend). DAPI was used as live stain.

#### Mouse cell culture

Cells were cultured in α-MEM with 10% FBS and 1% penicillin–streptomycin (Thermo Fisher Scientific; 15140-122). For in vitro osteogenic and chondrogenic differentiation assays, 12,000 FACS-purified mouse SSCs were cultured in expansion in wells of a 24-well tissue culture plate. Cells were washed in PBS, trypsinized and transferred to osteogenic differentiation medium containing 10% FBS, 100 μg/ml ascorbic acid and 10 mM β-glycerophosphate in α-MEM for 14 days. Alternatively, for chondrogenic differentiation micromass cultures were generated by a 5-μl droplet of cell suspension with approximately 1.5 × 10^7^ cells per ml pipetted in the center of a 24-well plate and cultured for 2 h in the incubator before adding warm chondrogenic medium consisting of α-MEM (high glucose) with 10% FBS, 100 nM dexamethasone, 1 μM L-ascorbic acid-2-phosphate and 10 ng/ml TGF-β1 (Invitrogen, cat#PHG9204). The micromass was maintained for 21 days with media changes every other day. At the end of differentiation cells were washed in PBS, fixed in 4% PFA and stained with Alizarin Red S (cat#A5533-25G, Sigma-Aldrich) solution (osteogenesis) or Alcian Blue (cat#A3157, Sigma-Aldrich) solution (chondrogenesis). For osteogenic potential Alizarin Red S stain was dissolved in 300 µl of 20% Methanol/10% Acetic Acid solution. After complete liberation of staining, 80 µl of each well was transferred into a new 96-well plate in duplicates and absorbance was measured at 450 nm. Chondrogenic potential was assessed by spectrophotometrically measuring absorption at 595 nm. For osteoclastogenesis assays flushed bone marrow cells from long bones were plated in 24-well plates at a density of 200,000 cells per well with α-MEM without phenol red, 1% GlutaMAX supplement (cat#35050061, Gibco), 10% FBS, 1% Penicillin-Streptomycin 10,000 U/ml, 1 µM prostaglandin E2 (cat#P0409, Sigma), and 10 ng/ml Csf1 recombinant murine protein (cat#315-02, Peprotech) for 3 days. Starting on day three, the media was changed daily to also include 10 ng/ml recombinant mouse RANKL (cat#315-11, Peprotech). Osteoclast culture continued for 10 days until large, multinucleated osteoclasts appeared. Plates were stained for osteoclasts using the TRAP kit (cat#387A, Sigma-Aldrich).

#### Human SSC (hSSC) culture

Freshly sorted, primary hSSCs were cultured in α-MEM with 10% human platelet derived lysate (HPL; cat#06962, STEMCELLtechnologies), 1% Penicillin-Streptomycin, 0.01% heparin and maintained at 37°C with 5% CO_2_. For in vitro differentiation assays hSSCs were treated and supplemented with the same osteogenic and chondrogenic differentiation protocols as described for mouse cells above. For viral overexpression experiments the coding region of Basigin (269aa) was cloned into a pCCLc backbone for manufacture of second generation lentivirus using Lenti-X 293T as packaging cells as previously described ^29^. The viral titer was determined functionally, based on amount of virus necessary to reach 90-95% eGFP expression. Cells were transduced using Lipofectamin3000 (cat#L3000001, ThermoFisher). For rescue experiments media was either supplemented with hPTH 1-34 by adding a final concentration of 2.5 nM 6h before each media change (to resemble intermittent exposure) or with 1 µg/mL of monoclonal Basigin antibody (cat# NB55-430, Novus Biologicals) during media change. Reactive oxygen species measurement was performed according to manufacturer’s instructions (cat# ab113851, Abcam).

#### Cell line experiments

Low passage VeraVec endothelial cells were thawed in a 10 cm dish with 10 mL of EGM-2 medium (cat#CC-3156, Lonza Bioscience). VeraVecs were grown at 37°C with 5% CO_2_ until confluent with media changes every 3 days. Before the assays, VeraVec culture was starved with low serum medium. For migration (wound scratch assay) ^30^, VeraVecs were lifted with 0.05% trypsin, spun down and resuspended in supernatant from 48h cultured SSCs. They were then plated at 1.5×10^5^ cells/well in a 24-well plate. To induce a gap for the migration, a scratch assay was generated using a 1000 µL micropipette tip, scratching vertically from one side of the well to the other. The gap was imaged, and area measured at times 0h and 12h. The area healed was calculated by (area at 0h - area at 12h)/(area at 0h) x 100%. For tube formation a matrix (cat#A1413201, Geltrex, ThermoFisher,) was deposited in wells of 48-well plate at an amount of 50 uL/cm^2^, for a total of 50 µL per well. As for migration, VeraVecs were lifted and resuspended in supernatant from SSCs. VeraVecs were plated at 1.5×10^4^ cells/well. Cells were imaged at 12h.

#### Subcutaneous xenografts

Primary hSSCs were transplanted into the dorsum of 3-month-old immunodeficient NSG mice (NOD.Cg-Prkdcscid Il2rgtm1Wjl/SzJ; JAX: 005557) as described previously ^31^. Briefly, freshly sorted patient-derived hSSCs were sorted, expanded to confluency and virally transduced to either overexpress Basigin or control. 1×10^6^ cells were mixed with 5 µl Matrigel and seeded on macroporous composite scaffolds formed of hydroxyapatite (HA) and poly(lactide-co-glycolide) (PLG) (HA-PLG) on ice. Scaffolds were fabricated using a gas foaming/particulate leaching method as previously described^32^. Microspheres composed of PLG (85:15; DLG 7E; Lakeshore Biomaterials, Birmingham, AL) were prepared using a double-emulsion process and lyophilized. Lyophilized microspheres (7.1 mg) were combined with 17.8 mg of synthetic HA (particle size, <200 nm; Aldrich Chemistry, St. Louis, MO) and 134.9 mg of NaCl (300 to 500 μm in diameter) to yield a 2.5:1:19 mass ratio of ceramic:polymer:salt. The powdered mixture was compressed under 2 metric tons for 1 min to form solid disks (final dimensions, 8 mm in diameter and 1.5 mm in height) using a Carver Press (Carver Inc., Wabash, IN). Compressed disks were exposed to high-pressure CO_2_ gas (5.5 MPa) for at least 24 hours, followed by rapid pressure release to prompt polymer fusion. Salt particles were leached from scaffolds in distilled H_2_O for 24 h to generate HA/PLG composite scaffolds. HA/PLG composite scaffolds were cut with a biopsy punch to produce scaffolds with final dimensions of 4 mm in diameter and 1.5 mm in height. Scaffolds were sterilized in 70% ethanol in 24 well plates for 20 minutes, followed by two rinses in sterile PBS. Sterile scaffolds were dried and kept until use. To promote cell adhesion, scaffolds were incubated in culture media at 37°C with 5% CO_2_ for 30 min directly before cell seeding. A small skin incision was made in the dorsum of anesthetized NSG mice and the cell containing scaffold was slid under the skin. Interrupted sutures were applied to close the incision and transplants were allowed to engraft and grow for 6 weeks.

#### Renal capsule transplantation

Renal capsule transplantations were conducted as previously described^10,12^. Briefly, in the anaesthetized mouse a 5-mm dorsal incision was made, and the kidney was exposed manually. Then, a 2-mm incision was created in the renal capsule using a needle bevel, and 5,000 FACS-purified mouse SSCs resuspended in 2 μl of Matrigel were transplanted beneath the capsule. The kidney was re-approximated manually and incisions were closed using sutures and staples. At time of surgery, mice were randomly divided into three groups. Mice either received subcutaneous placebo or Methylprednisolone pellets as described above. One group additionally was treated with hPTH (see above). Grafts were collected after 21 days.

#### Single cell RNA-sequencing

Femurs were collected from all the treatment groups, and each were processed, digested and prepared for FACS as described above. Single cell solutions of each treatment group were then pooled (*n* = 5 per group) and 1 × 10^6^ PI-Ter119-cells were sorted into collection tubes containing FACS buffer. Cells were then processed with 10X Chromium Next GEM Single Cell 3’ GEM kit (10X Genomics, v.3.1) according to the manufacturer’s instruction to target 5,000 cells per group. Barcoded samples were demultiplexed, aligned to the mouse genome (GRCm39.104), and UMI-collapsed with the Cell Ranger toolkit with standard settings (v.7.1.0, 10X Genomics), and sequenced on a partial lane Illumina NovaSeq platform. We used the Scanpy package (v.1.9.1.) to explore the data. First, quality filtering to exclude multiplets and cells of poor quality was conducted by only keeping cells with a gene count of more than 250 and fewer than 3,000, with less than 15% mitochondrial and 15% ribosomal gene content, leaving 8,614 cells. Genes expressed in fewer than three cells across all cells were also removed from downstream analysis. Data were log-normalized, cell cycle-regressed and scaled for analysis. Dimensionality reduction and Leiden clustering as well as subclustering were conducted choosing parameters based on PCA elbow plots. For single cell RNA-sequencing of SSC-derived grafts the tissue the tissue was dissected out and processed as described for femur bone processing to isolate SSCs above. Single cell solution was then stained with PI and Ter119, and living (PI-negative), non-red blood cells (Ter119-negative) were sorted into FACS buffer for each group. Based on yield, cells were then processed with 10X Chromium Next GEM Single Cell 3’ GEM kit (10X Genomics, v.3.1) according to the manufacturer’s instruction to target 2,000 cells for Placebo and GC+PTH groups as well as 500 cells for GC group. All other steps were conducted as described above. A total of 595 graft-derived cells across all groups passed stringent quality filtering. Data are available under GEO accession number GSE253044.

#### Statistical analysis

Statistical significance between placebo and treatment groups was determined using two-tailed, unpaired Student’s t-test or One-way ANOVA for multiple groups comparison with LSD Fisher test unless stated otherwise in the figure legend (GraphPad Prism; version 10). Data were tested for normality by Shapiro-Wilk test. Statistical significance was defined as p < 0.05. All data points refer to biological replicates and are presented as mean ± standard error of the mean (SEM) unless otherwise stated in figure legend.

## Data Availability

Single cell RNA-sequencing data was deposited in the GEO database with access number GSE253044.

## AUTHOR CONTRIBUTIONS

N.L. conceived the work, supervised the study and wrote the manuscript. T.A. co-conceived the study, designed experiments, performed experiments, co-supervised the study and wrote the manuscript. C.C. co-supervised the study and helped design experiments. D.M. and K.C. performed the majority of the experiments and analyses. E.H., K.W., A.M., F.C., Y.W., M.M., E.W., A.C., A.S., K.L., F.F. contributed expertise, performed experiments and/or data analysis.

## ACKNOWLEDGMENTS

We thank Stanford University Institute for Stem Cell Biology and Regenerative Medicine FACS core (NIH S10 RR02933801) for experimental support; and NIH S10 1S10OD028493-01A1 (Principal Investigator: Charles.K.F. Chan). MicroCT work on the Bruker Skyscan 1276 was supported by NIH S10 Shared Instrumentation Grant (1S10OD02349701, PI Timothy C. Doyle). Further funding for this work was provided by NIH–NIA K99/R00AG066963 to T.A., NIH–NIA K99/R00AG049958-01A1, Stanford Wu Tsai Human Performance Alliance Funds, Siebel Foundation, the Heritage Medical Foundation, Prostate Cancer Foundation, the American Federation for Aging Research (AFAR)–Arthritis National Research Foundation (ANRF), a seed grant from the WHSDM Stanford Women’s Health and Sex Differences in Medicine Center, and an endowment from the DiGenova Family to C.C.. E.W. and K.W. were supported by the California Institute of Regenerative Medicine (CIRM) EDUC4-12792 Research Training Program.

## COMPETING FINANCIAL INTERESTS

The authors have declared that no conflict of interest exists.

**Extended Data Figure 1. Micro-CT bone parameters of experimental groups. (a)** Representative H&E staining images of metaphyseal regions of distal femurs of different experimental groups. **(b)** Quantification of trabecular bone thickness (Tb.Th), number (Tb.N) and spacing (Tb.Sp) shown as percentage change compared to day 0. **(c)** Representative microCT images of cortical bone at day-28 and day-56. **(d)** Quantification of femoral cortical thickness (Cort.Th) and cortical area (Cort.Ar.). All experiments n=6-9 mice per group. Statistical testing between Placebo and other group by unpaired student t-test. *p<0.05, **p<0.01 ***p<0.001, ****p<0.0001. Scale bars, 50 µm.

**Extended Data Figure 2. Bone parameters of experimental groups. (a)** Representative Calcein double labeling for each experimental group. **(b)** Quantification of mineral apposition rate (MAR) and bone formation rate (BFR) based on Calcein double labeling. n=4 mice per group. **(c)** Quantification of osteoclast surface per bone surface (OC.s/BS) by TRAP labeling. n=3 mice per group. **(d)** Quantification of bone marrow derived osteoclast formation from bones harvested at day-56 (left) and representative images thereof (right). n=5 mice per group. **(e)** Measurement of growth plate height based on H&E overview stain. n=3 mice per group. All data shown as mean ± SEM. Statistical testing between Placebo and other group by unpaired student t-test. *p<0.05, **p<0.01 ***p<0.001, ****p<0.0001. Scale bar, 50µm.

**Extended Data Figure 3. Flow cytometric gating strategies. (a)** Representative gating strategy for skeletal stem cell (SSC) and bone-cartilage-stromal-progenitor cell (BCSP) populations. **(b)** Flow cytometric based quantification of transient BCSP (CD45-Ter119-Tie2-CD90-6c3-CD105+CD51+) in femurs of experimental groups. n=3-8. **(c)** Representative gating strategy for CD31+ endothelial cell populations. **(d)** Representative immunohistochemistry staining for Endomucin (Emcn) in bone marrow of day-28 experimental groups. All data shown as mean ± SEM. Statistical testing between Placebo and other group by unpaired student t-test. *p<0.05, **p<0.01 ***p<0.001, ****p<0.0001. Scale bars, 20µm.

**Extended Data Figure 4. Cellular composition of bone marrow and blood in anti-Basigin treated mice. (a)** Representative gating strategy for hematopoietic stem and progenitor cells. **(b)** Frequency of adipogenic (APC, Lin-Sca1+CD24-) progenitor cells in bone marrow. n=5. **(c)** Myeloid cell fraction in mouse blood samples. n=5. **(d)** Flow cytometric analysis of hematopoietic stem and progenitor cells in bone marrow. n=5. CLP: common lymphoid progenitor, GMP: granulo-monocyte progenitor, CMP: common myeloid progenitor, MEP: myelo-erythroid progenitor, LSK: Lin^low^Sca1+cKit+ hematopoietic stem and progenitor cells, MPPa: multipotent progenitor a, MPPb: multipotent progenitor b, MPPc: multipotent progenitor c, pHSC: phenotypic hematopoietic stem cell, myHSC: myeloid skewed HSC, balHSC: balanced HSC. All data shown as mean ± SEM. Statistical testing by one-way ANOVA with Fisher-LSD test. *p<0.05, **p<0.01 ***p<0.001, ****p<0.0001.

**Extended Data Figure 5. Cellular composition of bone marrow in anti-Basigin treated aged mice. (a)** Representative immunohistochemistry of endothelial Endomucin expression (green) bone marrow of aged mice treated with control IgG or anti-Basigin. **(b)** Flow cytometric analysis of skeletal cell populations. **(c)** Flow cytometric analysis of myeloid and lymphoid cell frequency in blood samples. **(d)** Flow cytometric analysis of hematopoietic stem and progenitor cell populations in bone marrow. All data shown as mean ± SEM. Statistical testing between Placebo and other group by unpaired student t-test. *p<0.05, **p<0.01 ***p<0.001, ****p<0.0001.

