## Supplemental for "Basigin Links Altered Skeletal Stem Cell Lineage Dynamics with Glucocorticoid-induced Bone Loss and Impaired Angiogenesis"

### Extended Data Figure 1

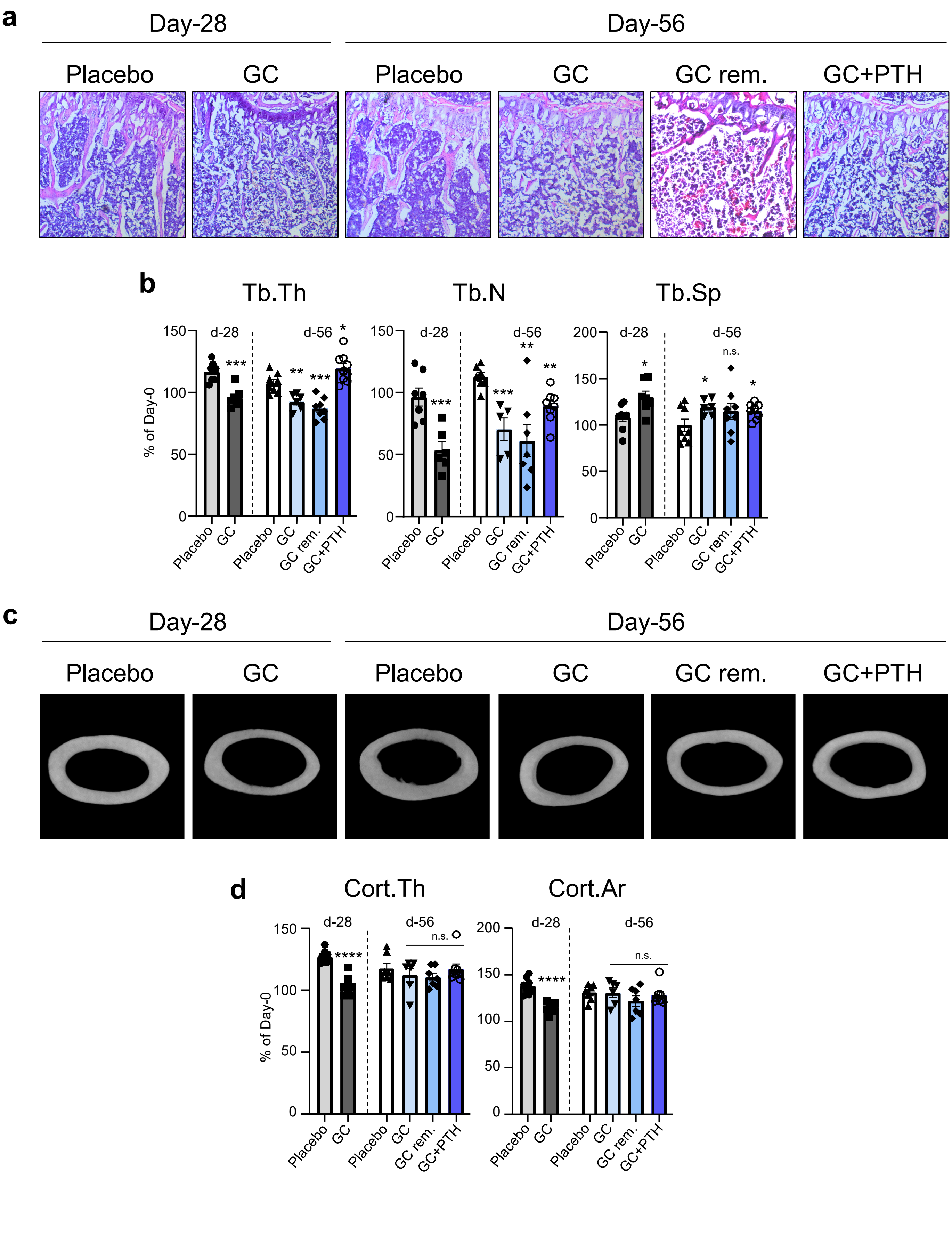


**Extended Data Figure 1. Micro-CT bone parameters of experimental groups. (a)** Representative H&E staining images of metaphyseal regions of distal femurs of different experimental groups. **(b)** Quantification of trabecular bone thickness (Tb.Th), number (Tb.N) and spacing (Tb.Sp) shown as percentage change compared to day 0. **(c)** Representative microCT images of cortical bone at day-28 and day-56. **(d)** Quantification of femoral cortical thickness (Cort.Th) and cortical area (Cort.Ar.). All experiments n=6-9 mice per group. Statistical testing between Placebo and other group by unpaired student t-test. *p<0.05, **p<0.01 ***p<0.001, ****p<0.0001. Scale bars, 50 μm.

### Extended Data Figure 2

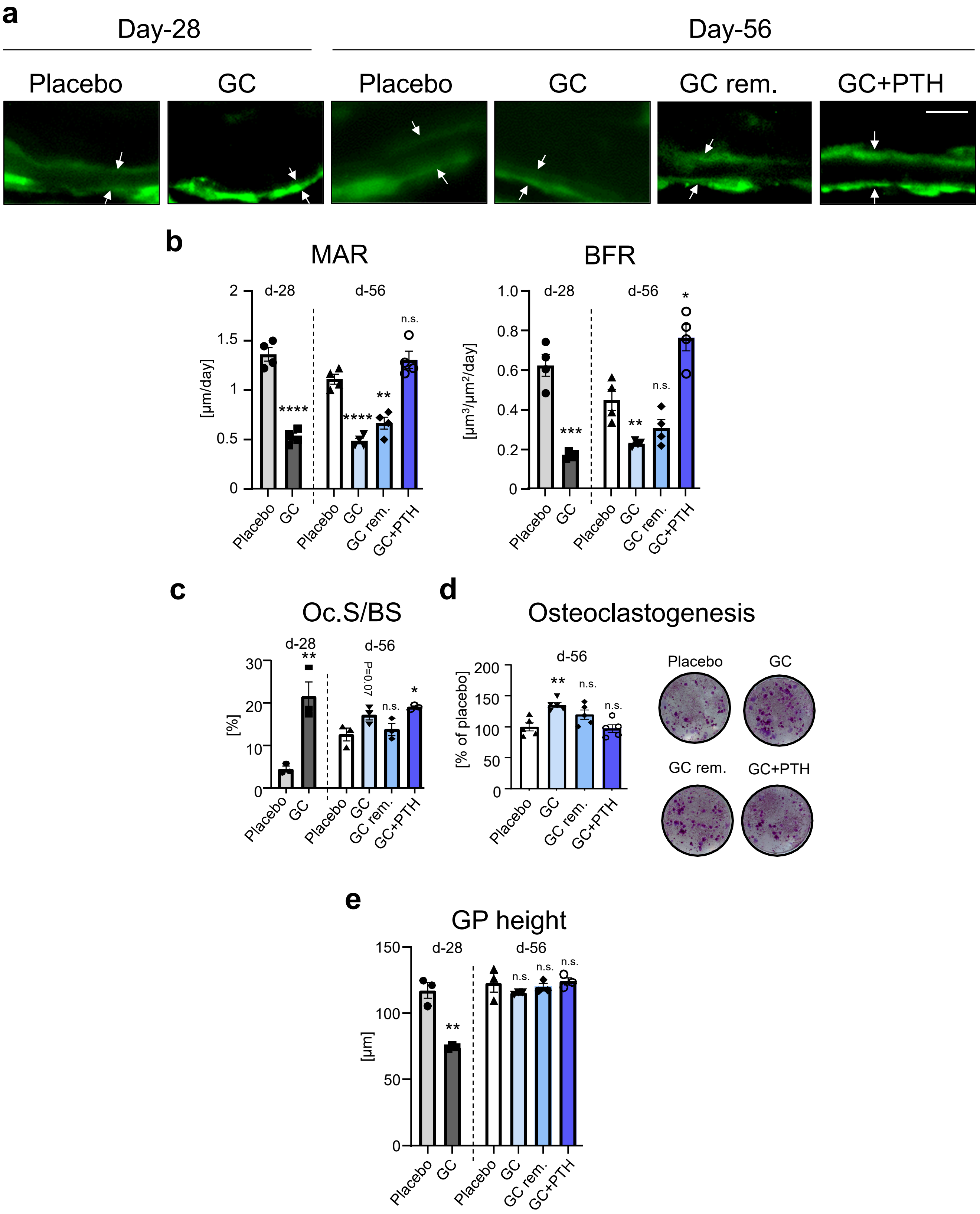


**Extended Data Figure 2. Bone parameters of experimental groups. (a)** Representative Calcein double labeling for each experimental group. **(b)** Quantification of mineral apposition rate (MAR) and bone formation rate (BFR) based on Calcein double labeling. n=4 mice per group. **(c)** Quantification of osteoclast surface per bone surface (OC.s/BS) by TRAP labeling. n=3 mice per group. **(d)** Quantification of bone marrow derived osteoclast formation from bones harvested at day-56 (left) and representative images thereof (right). n=5 mice per group. **(e)** Measurement of growth plate height based on H&E overview stain. n=3 mice per group. All data shown as mean ± SEM. Statistical testing between Placebo and other group by unpaired student t-test. *p<0.05, **p<0.01 ***p<0.001, ****p<0.0001. Scale bar, 50µm.

### Extended Data Figure 3

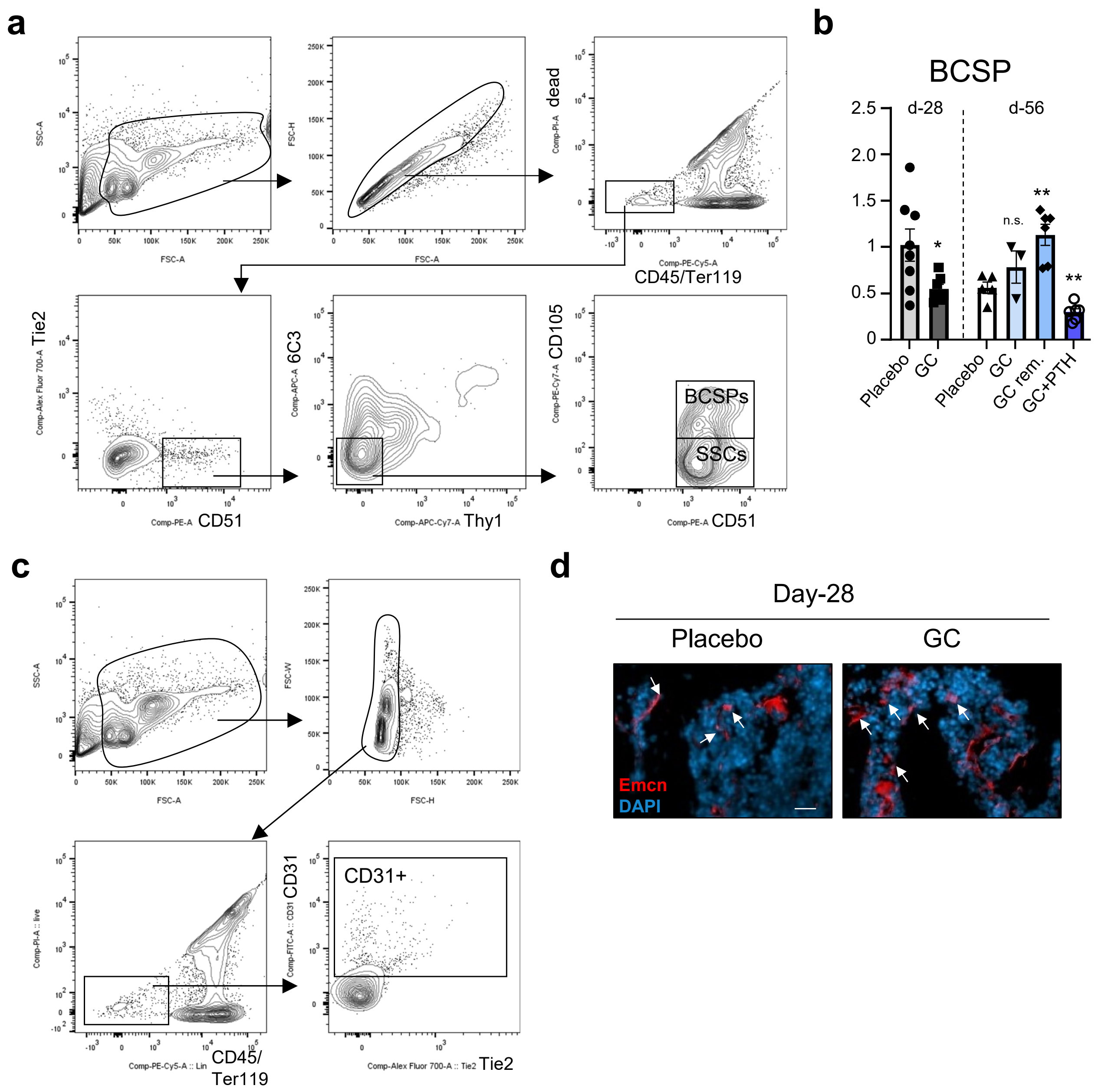


**Extended Data Figure 3. Flow cytometric gating strategies. (a)** Representative gating strategy for skeletal stem cell (SSC) and bone-cartilage-stromal-progenitor cell (BCSP) populations. **(b)** Flow cytometric based quantification of transient BCSP (CD45-Ter119-Tie2-CD90-6c3-CD105+CD51+) in femurs of experimental groups. n=3-8. **(c)** Representative gating strategy for CD31+ endothelial cell populations. **(d)** Representative immunohistochemistry staining for Endomucin (Emcn) in bone marrow of day-28 experimental groups. All data shown as mean ± SEM. Statistical testing between Placebo and other group by unpaired student t-test. *p<0.05, **p<0.01 ***p<0.001, ****p<0.0001. Scale bars, 20μm.

### Extended Data Figure 4

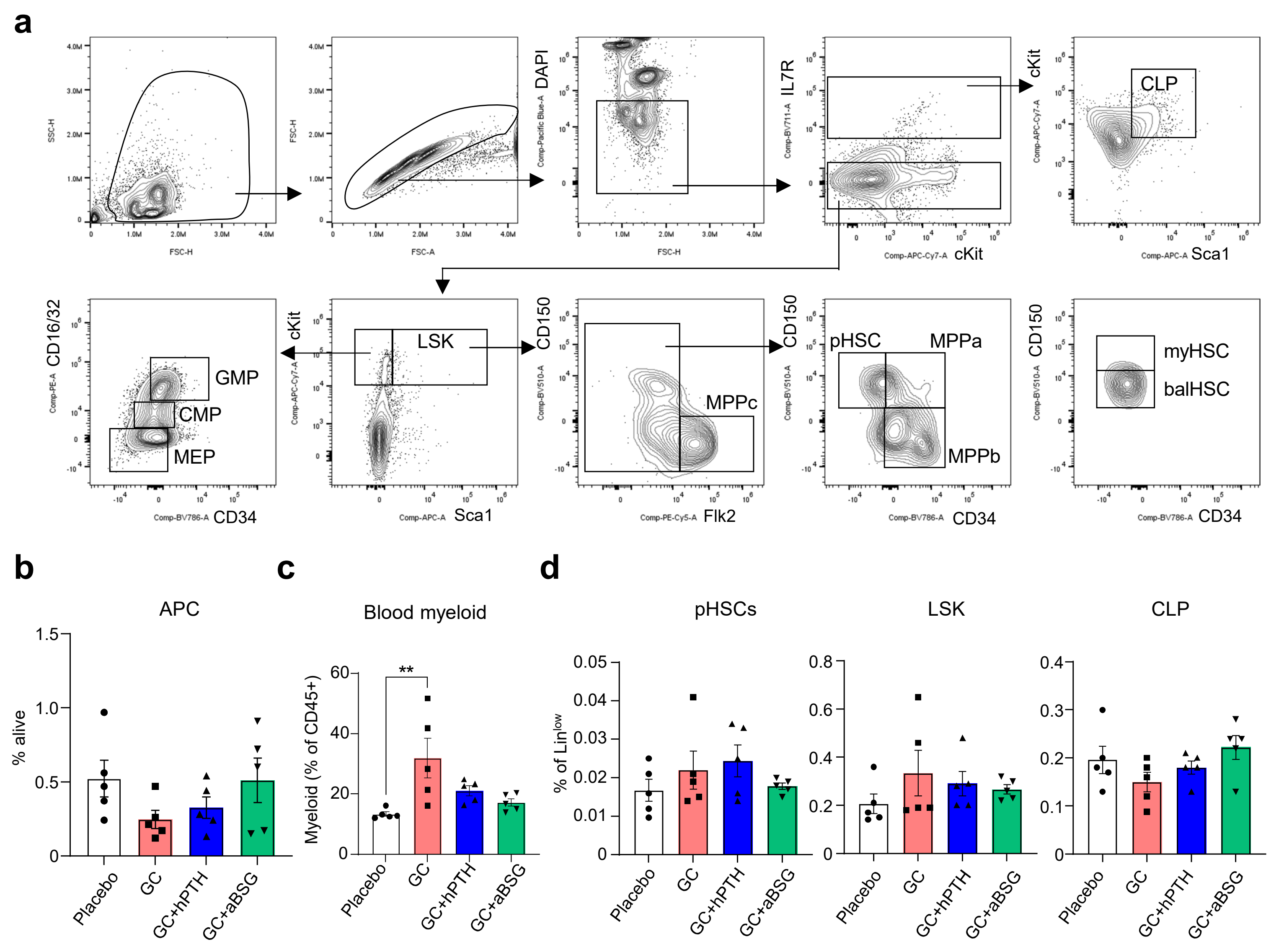


**Extended Data Figure 4. Cellular composition of bone marrow and blood in anti-Basigin treated mice. (a)** Representative gating strategy for hematopoietic stem and progenitor cells. **(b)** Frequency of adipogenic (APC, Lin-Sca1+CD24-) progenitor cells in bone marrow. n=5. **(c)** Myeloid cell fraction in mouse blood samples. n=5. **(d)** Flow cytometric analysis of hematopoietic stem and progenitor cells in bone marrow. n=5. CLP: common lymphoid progenitor, GMP: granulo-monocyte progenitor, CMP: common myeloid progenitor, MEP: myelo-erythroid progenitor, LSK: $Lin^{low}$Sca1+cKit+ hematopoietic stem and progenitor cells, MPPa: multipotent progenitor a, MPPb: multipotent progenitor b, MPPc: multipotent progenitor c, pHSC: phenotypic hematopoietic stem cell, myHSC: myeloid skewed HSC, balHSC: balanced HSC. All data shown as mean ± SEM. Statistical testing by one-way ANOVA with Fisher-LSD test. *p<0.05, **p<0.01 ***p<0.001, ****p<0.0001.

### Extended Data Figure 5

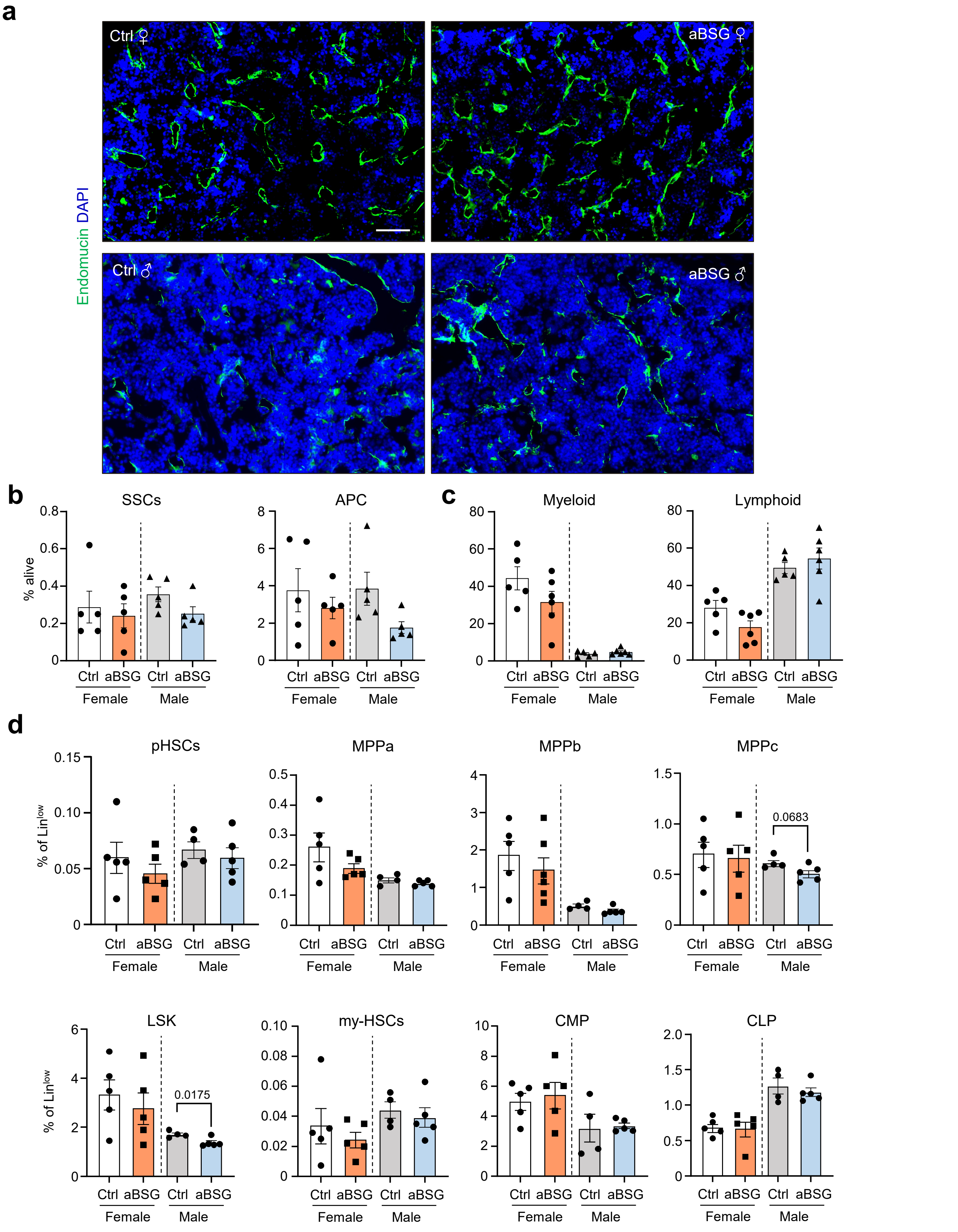


**Extended Data Figure 5. Cellular composition of bone marrow in anti-Basigin treated aged mice. (a)** Representative immunohistochemistry of endothelial Endomucin expression (green) bone marrow of aged mice treated with control IgG or anti-Basigin. **(b)** Flow cytometric analysis of skeletal cell populations. **(c)** Flow cytometric analysis of myeloid and lymphoid cell frequency in blood samples. **(d)** Flow cytometric analysis of hematopoietic stem and progenitor cell populations in bone marrow. All data shown as mean ± SEM. Statistical testing between Placebo and other group by unpaired student t-test. *p<0.05, **p<0.01 ***p<0.001, ****p<0.0001. Scale bar, 100µm.
